# Dietary Protein Source Shapes Gut Microbial Structure and Predicted Function: A Meta-Analysis with Machine Learning

**DOI:** 10.1101/2025.04.23.650130

**Authors:** Samson Adejumo, Garry Lewis, Pritha Das, Casey Kin Yun Lim, Judy Malas, Angus Nnamdi Oli, Jacob M. Allen, Jarrad Hampton-Marcell

## Abstract

**Background:** Dietary proteins shape gut microbial ecology, yet the taxonomic and functional consequences of plant- versus animal-based proteins remain poorly defined. Although digestibility and fermentation profiles differ by protein type, a systematic evaluation of how these differences influence microbial diversity, community structure, and metabolic capacity is lacking. This meta-analysis integrates taxonomic, machine learning, and functional inference approaches to identify microbial and metabolic signatures associated with dietary protein source in murine models.

**Methods:** Following PRISMA guidelines, we analyzed 16S rRNA sequencing data from 10 murine studies (n = 187) comparing plant- and animal-protein diets. Alpha diversity was assessed using Shannon, Inverse Simpson, and Chao1 indices, and beta diversity with Aitchison distances. Differentially abundant taxa were identified using LEfSe and class-weighted Random Forest models. Functional potential was inferred with PICRUSt2, and taxon–pathway relationships were explored using correlation and network analyses.

**Results:** Plant-protein diets increased gut microbial diversity across all alpha diversity metrics and were associated with higher representation of saccharolytic and nitrogen-recycling genera such as Bacteroides, Muribaculaceae, and Allobaculum. Animal-protein diets favored proteolytic taxa, including Clostridium sensu stricto 1 and Colidextribacter. Microbial community structure differed significantly between diets (ANOSIM R = 0.663, p < 0.001). Random Forest models achieved >88% accuracy (AUC = 0.995) in predicting dietary groups, and LEfSe identified consistent discriminating taxa. Functional profiling showed that plant-based diets enriched pathways linked to short-chain fatty acid and aromatic amino acid metabolism, whereas animal-based diets favored sulfur- and branched-chain amino acid-associated pathways. Network analysis identified Muribaculaceae as a plant-associated hub and Lactobacillus as an animal-associated hub.

**Conclusion:** Dietary protein source significantly influences gut microbiota composition and functional potential in mice. Plant- and animal-based proteins generate distinct metabolic signatures with implications for nitrogen cycling, sulfur metabolism, and microbial ecology. Future controlled dietary studies that harmonize protein source with other macronutrient variables are needed to isolate protein-specific effects.

**Statement of Significance:** This study presents the first standardized meta-analysis of murine protein-intervention microbiome datasets, integrating taxonomic, machine learning, and predicted functional profiling to identify robust microbial and metabolic signatures that differentiate plant- from animal-based protein diets. By harmonizing evidence across diverse studies, it clarifies foundational ecological responses to protein source and provides mechanistic hypotheses that can inform future controlled nutrition and microbiome research

## 1.0 Introduction

The gut microbiome is a complex and dynamic ecosystem that engages in bidirectional interactions with host nutrition and immunity and plays a critical role in metabolic health (Wu *et al.*, 2011). While the effects of dietary carbohydrates, particularly in promoting saccharolytic fermentation and short-chain fatty acid (SCFA) production, are well established, the influence of dietary protein on gut microbial ecology is less clearly defined (Rowland *et al.*, 2018). Dietary proteins that escape digestion in the upper gastrointestinal tract enter the colon, where they undergo fermentation by resident microbes. The outcomes of this process vary by protein source and depend on the compositional and functional capacities of the gut microbiota (Scott *et al.*, 2013; Beaumont *et al.*, 2017).

Microbial protein metabolism can proceed through saccharolytic- or proteolytic-associated pathways. Saccharolytic fermentation of amino acids can generate SCFAs that support epithelial barrier integrity and modulate host immunity (Canfora *et al.*, 2019). In contrast, proteolytic degradation produces nitrogenous compounds such as ammonia, hydrogen sulfide, and phenolic metabolites. These byproducts have been associated with microbial imbalance, epithelial damage, and inflammatory responses (Windey *et al.*, 2012; Rooks & Garrett, 2016). Together, these distinct metabolic routes suggest that the net impact of protein fermentation is shaped by a combination of substrate availability and microbial community composition, and the metabolic specialization of resident taxa (Adejumo *et al.*, 2023).

Importantly, not all protein sources influence the microbiota equally. Plant proteins are often accompanied by fiber and polyphenols, which foster saccharolytic fermentation and SCFA production, contributing to anti-inflammatory effects and gut barrier reinforcement (den Besten *et al., 2013*; Vinelli *et al.*, 2022). However, plant proteins may also be less digestible and lower in certain essential amino acids, potentially limiting their nutritional completeness (Scott *et al.*, 2013; Wang *et al.*, 2020; Rutherfurd *et al.*, 2015). In contrast, animal proteins provide a more complete amino acid profile, beneficial for muscle synthesis and maintenance, particularly in aging populations (Phillips *et al.*, 2016) but are typically higher in sulfur-containing amino acids like methionine and cysteine. These amino acids can fuel proteolytic fermentation and lead to elevated levels of hydrogen sulfide and ammonia, which have been associated with mucosal inflammation and impaired gut barrier function (Kitada *et al.*, 2019; Ling *et al.*, 2023).

Despite growing interest, relatively few studies have systematically compared the microbial responses to plant- versus animal-based proteins, particularly at both the taxonomic and functional levels. Many investigations focus on a single protein type or infer function without linking it to compositional shifts. This limits our ability to identify microbial biomarkers of protein source or to understand how these shifts contribute to gut health and disease.

Murine models remain essential for addressing these questions because they enable precise manipulation of dietary protein source while controlling for genetic background, environmental exposures, and non-protein dietary components—conditions that are not feasible in free-living human populations (Hildebrand *et al.*, 2013; Nguyen *et al.*, 2015; Turnbaugh *et al.*, 2009). Such experimental consistency allows for mechanistic insights into how protein-specific perturbations structure microbial communities and their functional outputs.

To address this gap, we conducted a meta-analysis of 16S rRNA gene amplicon sequencing data from ten murine studies (n = 187), integrating statistical, machine learning, and predictive functional profiling approaches. We assessed microbial alpha and beta diversity, taxonomic composition, and classification accuracy using Random Forest and LEfSe. Functional inference via PICRUSt2 was used to identify enriched metabolic pathways, while correlation and network analyses explored taxa–function relationships. We report that plant protein consumption is associated with greater microbial diversity and enrichment of taxa involved in SCFA production and nitrogen recycling. Conversely, animal protein favors the enrichment of proteolytic taxa linked to ammonia, sulfur, and phenolic metabolite production. Together, these findings offer a taxonomic and functional framework for understanding how dietary protein source modulates the gut microbiome, and they provide a foundation for future mechanistic studies linking diet, microbial metabolism, and host health.

## 2.0 Materials and Methods

### 2.1 Search Strategy

This meta-analysis followed the Preferred Reporting Items for Systematic Review and Meta-Analysis (PRISMA) guidelines (Page *et al.*, 2021) to ensure a transparent, systematic, and reproducible study selection process. PRISMA provides a structured framework for identifying, screening, and selecting studies in a manner that minimizes bias and enhances the reliability of findings. A comprehensive literature search was conducted to identify murine studies published in different databases from inception to March 31, 2024. Peer-reviewed, English-written articles were retrieved from (1) Web of Science, (2) PubMed, (3) MEDLINE Complete, (4) Scopus, (5) Google Scholar, and (6) manual searching. The Boolean search string used was: *(“protein fermentation” OR “protein digestion” OR “protein intake” OR “dietary protein” OR “protein metabolism” OR “protein consumption” OR “protein diet” ) AND ( “microbiome” OR “microflora” OR “microbial” OR “commensal” OR “microbiota” OR “bacteria” ) AND ( “gut” OR “intestinal” OR “gastrointestinal” OR “intestine” OR “digestive tract” OR “GIT” OR “enteric” OR “ileum” OR “colon” OR “colonic” OR “duodenum” OR “jejunum” OR “fecal” OR “faecal” ) AND ( “mice” OR “rat” OR “rats” OR “mouse” OR “rodent” OR “rodents” OR “murine” )*.

### 2.2 Selection Criteria

The study selection process followed PRISMA guidelines and consisted of four phases: (1) Identification of relevant records, (2) Screening of abstracts for eligibility, (3) Full-text eligibility assessment, and (4) Final inclusion in the meta-analysis. The PRISMA flow diagram (**Figure 1**) summarizes the number of records identified, screened, excluded, and retained for analysis. Studies were included if they met the following criteria: (1) involved murine models (mice or rats), (2) examined dietary protein interventions, (3) utilized 16S rRNA gene sequencing, (4) had clearly distinguishable treatment and control groups, and (5) provided gut microbiota composition data. Studies were excluded if they were editorials, letters, reviews, comments, or other meta-analyses, lacked a protein intervention, involved non-murine or in vitro studies, or had unpublished data or unclear dietary group differentiation. Search results were exported to Mendeley for initial deduplication, followed by a second deduplication in Zotero. The remaining records were manually screened by title and abstract to exclude studies irrelevant to the research objectives.

**Figure 1.**
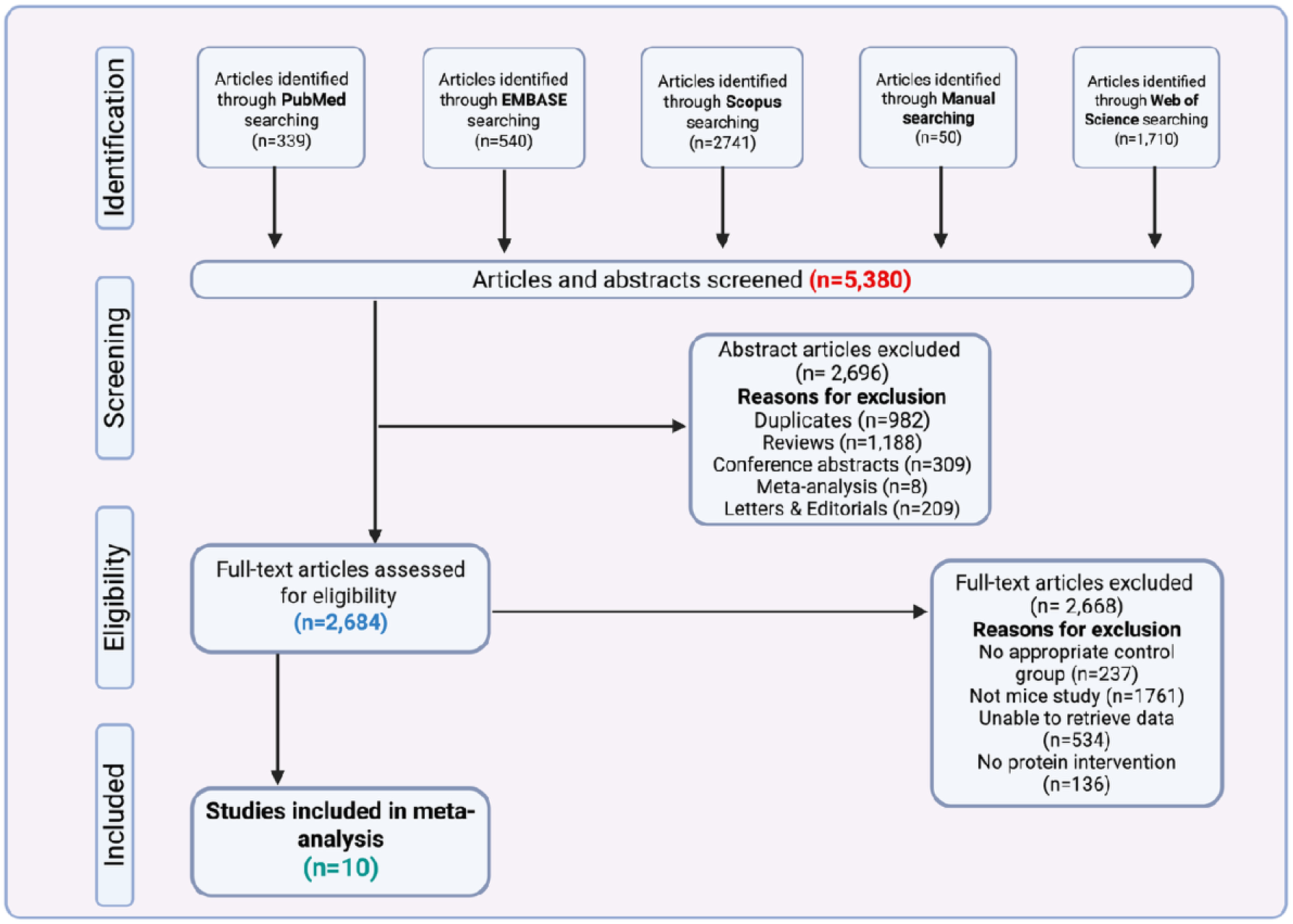
PRISMA Flow Diagram for Study Selection summarizing the systematic search and screening process for identifying eligible studies included in the meta-analysis.

### 2.3 Bioinformatics and Data Processing

Raw sequencing data (FASTQ files) were retrieved from the NCBI and ENA repositories using SRA accession numbers reported in the included studies. Amplicon sequence processing was conducted using QIIME2-amplicon-2024.5 (Bolyen *et al.*, 2019; Ziemski *et al.*, 2022). Reads were demultiplexed and quality-filtered using the q2-demux plugin, followed by denoising with DADA2 (v1.28.0) to generate high-resolution amplicon sequence variants (ASVs) (Callahan *et al.*, 2016). Taxonomic assignment was performed against the SILVA-138 reference database using a Naïve Bayes classifier implemented via the q2-feature-classifier plugin (Bokulich *et al.*, 2018). ASVs with confidence scores below 70% were discarded to improve taxonomic reliability.

Processed data, including feature tables, phylogenetic trees, and taxonomic classifications, were imported into RStudio (v4.4.1) for downstream analyses. Microbiome data were integrated using the phyloseq package (v1.48.0), and statistical analyses were conducted using ggplot2 (v3.5.1), patchwork (v1.3.0), ggpubr (v0.6.0), pheatmap (v1.0.12), and dplyr (v1.1.4) (McMurdie & Holmes, 2012; Wen *et al.*, 2023).

Functional metagenomic prediction was performed using PICRUSt2 (v2.5.2), which infers Kyoto Encyclopedia of Genes and Genomes (KEGG) Orthologs (KOs) and MetaCyc pathways from 16S rRNA gene data via evolutionary modeling. The predicted KO and pathway tables were normalized to relative abundance and exported for downstream differential abundance testing and network analysis

Among the included studies, nine targeted the V3–V4 region of the 16S rRNA gene using either the 338F/806R or 515F/806R primer pairs. Both forward primers amplify the V3 region, but 338F (5′-ACTCCTACGGGAGGCAGCAG-3′) is known for its broad bacterial coverage in murine gut microbiota, while 515F (5′-GTGCCAGCMGCCGCGG-3′) is commonly used in high-throughput studies due to its compatibility with barcoding and multiplex sequencing (Parada *et al.*, 2016). The reverse primer 806R (5′-GGACTACHVGGGTWTCTAAT-3′) was used consistently across all V3–V4 studies. One study amplified the V1–V2 region using 27F (5′-AGAGTTTGATCMTGGCTCAG-3′) and 338R (5′-TGCTGCCTCCCGTAGGAGT-3′), which target a more upstream and variable portion of the gene. Primer selection can influence sequencing depth, taxonomic bias, and classification accuracy (Klindworth *et al.*, 2013; Parada *et al.*, 2016), so all primer sequences and target regions are summarized in **Supplementary Table 1**

**Table 1:** Characteristics of studies included in the meta-analysis.

| Country, Year, First Author | Title | Bioproject accession | Protein source | Sample number | Age of mice (weeks) |
| --- | --- | --- | --- | --- | --- |
| Australia, 2022<br>Jian Tan | Dietary protein increases T-cell-independent sIgA production through changes in gut microbiota-derived extracellular vesicles | PRJEB39583 | Casein | 8 | 6 |
| Japan, 2020<br>Hiroaki Masuoka | The influences of low protein diet on the intestinal microbiota of mice | PRJDB8898 | Casein | 30 | 9 |
| China, 2022<br>Yantao Yin | Dietary oxidized beef protein alters gut microbiota and induces colonic inflammatory damage in C57BL/6 mice | PRJNA872483 | Oxidized Beef | 24 | 5 |
| China, 2022<br>Yifan Zhu | Reno-Protective Effect of Low Protein Diet Supplemented With α-Ketoacid Through Gut Microbiota and Fecal Metabolism in 5/6 Nephrectomized Mice | PRJNA810785 | Low Protein diet | 10 | 8 |
| China, 2020 | Processing Method Altered | PRJNA607338 | Casein, | 30 | 4 |
| **Yunting Xie** | Mouse Intestinal Morphology and Microbial Composition by Affecting Digestion of Meat Proteins |  | Pork, Sausage, Soy protein |  |  |
| **Canada, 2021**<br>**Béatrice S.-Y. Choi** | Feeding diversified protein sources exacerbates hepatic insulin resistance via increased gut microbial branched-chain fatty acids and mTORC1 signaling in obese mice | PRJEB37442 | Casein | 12 | 10 |
| **China, 2020**<br>**Muhammad Umair Ijaz** | Meat Protein in High-Fat Diet Induces Adipogenesis and Dyslipidemia by Altering Gut Microbiota and Endocannabinoid Dysregulation in the Adipose Tissue of Mice | PRJNA548036 | Beef, Casein, Chicken, Pork | 27 | 7 |
| **China, 2022**<br>**Wuling Zhong** | High-protein diet prevents fat mass increase after dieting by counteracting Lactobacillus-enhanced lipid absorption | PRJNA868153 | Casein | 14 | 8 |
| **China, 2022**<br>**Ranran Zhang** | Oral administration of branched-chain amino acids ameliorates high-fat diet-induced metabolic-associated fatty liver disease via gut microbiota-associated mechanisms | PRJNA852193 | Branched-chain amino acids | 12 | 6 |
| **Italy, 2020**<br>**Chiara Ruocco** | Manipulation of Dietary Amino Acids Prevents and Reverses Obesity in Mice Through Multiple Mechanisms That Modulate Energy Homeostasis | PRJEB25686 | Essential amino acids, Casein | 20 | 8 |

### 2.4 Statistical Analysis

#### 2.4.1 Data Transformation

Centered log-ratio (CLR) transformation was applied to genus-level relative abundance tables before all compositional analyses. CLR normalization addresses the inherent compositional constraints of microbiome data by expressing abundances as log-ratios, thereby reducing spurious correlations and enabling valid Euclidean distance–based comparisons (Gloor *et al.*, 2017; Quinn *et al.*, 2018). All CLR-based outputs in this study, including Aitchison distances and enrichment estimates, were generated using the microbiome and phyloseq packages in R.

#### 2.4.2 Handling of Missing Data

To ensure consistency across datasets, microbiome features with zero counts were retained as biological zeros rather than treated as missing values. Samples lacking essential metadata (e.g., protein level, animal age, country of origin, or sample isolation site) were excluded only from analyses that required those variables as fixed or random effects but were retained in descriptive summaries when appropriate. For functional pathway data, features with >20% missing values across studies were removed prior to analysis, and remaining missing entries were imputed as zeros, consistent with conventions in compositional microbiome analysis (Gloor *et al.*, 2017). These steps minimized bias arising from incomplete metadata while preserving the integrity of taxonomic and functional profiles across studies.

#### 2.4.3 Alpha and Beta Diversity

Alpha diversity indices (Shannon, Simpson, Chao1) were used to assess microbial richness and diversity (Li *et al.*, 2022; Zaheer *et al.*, 2018; Deng *et al.*, 2022). Comparisons between plant- and animal-protein diets were performed using the Wilcoxon Rank-Sum test. Beta diversity was assessed using Aitchison distance (Euclidean distance on centered log-ratio [CLR]–transformed data) and visualized via Principal Component Analysis (PCA). Analysis of Similarities (ANOSIM) tested whether microbial composition varied more between dietary groups than within groups (Samuthpongtorn *et al.*, 2022). Permutational analysis of variance (PERMANOVA) was conducted to quantify the variance in microbial composition explained by dietary protein sources (Tang *et al.*, 2016).

#### 2.4.4 Generalized Linear Mixed Model (GLMM)

To account for inter-study heterogeneity and sampling variability, a Generalized Linear Mixed Model (GLMM) was implemented using the lme4 (v1.1-34) package in R. Shannon diversity served as the continuous outcome variable, enabling estimation of the independent association between dietary protein source and alpha diversity. Dietary protein source and protein concentration were modeled as fixed effects, while random intercepts were specified for country of origin, sample isolation site (fecal, colonic, ileal, small intestine), and animal age (in weeks) to control for study-level and anatomical variation. Different model configurations were evaluated, and final model selection was guided by goodness-of-fit statistics, including Akaike Information Criterion (AIC) and Bayesian Information Criterion (BIC). This modeling approach enabled robust estimation of microbial diversity metrics while accounting for clustering and potential confounding across datasets (Madi *et al.*, 2023; Sweeny *et al.*, 2023).

#### 2.4.5 Machine Learning Classification and Microbial Feature Selection

To classify dietary protein source and identify microbial biomarkers, a supervised Random Forest (RF) model was implemented using the randomForest package in R (Manandhar *et al.*, 2021). Three RF approaches were compared: (1) unadjusted RF, trained on the original imbalanced dataset; (2) class-weighted RF, which accounted for class imbalance by assigning inverse frequency weights to underrepresented groups; and (3) synthetic sampling RF, using the ROSE (Random OverSampling Examples) technique to balance the dataset.

Microbial relative abundance data at the genus level were used as input features. The dataset was randomly partitioned into training (70%) and testing (30%) sets. Model performance was evaluated using the out-of-bag (OOB) error rate, confusion matrices, and area under the receiver operating characteristic curve (AUC). Feature importance was determined using the Mean Decrease in Accuracy (MDA), which ranked genera based on their contribution to classification performance

To complement the RF analysis, LEfSe (Linear Discriminant Analysis Effect Size) was performed using the microbiomeMarker package (v1.10.0) in R to independently identify microbial taxa significantly enriched in plant- versus animal-protein diets. LEfSe was applied to genus-level relative abundance data, using an LDA score threshold >3.0 and adjusted p-value <0.05. A taxonomic cladogram was generated to visualize differentially enriched microbial features between dietary groups (Segata *et al.*, 2011)

#### 2.4.6 Receiver Operating Characteristic (ROC) Analysis

Receiver Operating Characteristic (ROC) analysis was used to assess the predictive performance of the three Random Forest (RF) models. ROC curves were generated using the pROC package (v1.18.0) in R. Each model was trained to classify diets based on microbial genus-level profiles, distinguishing between plant and animal protein sources. Predictions from the test dataset were used to generate ROC curves, and the Area Under the Curve (AUC) was computed to quantify model performance. Higher AUC values indicated stronger sensitivity–specificity trade-offs and improved classification capability. Overall classification accuracy was also reported to compare performance across sampling strategies

ROC analysis is commonly used in microbiome studies to assess classification models, particularly in dietary and clinical interventions (Zhou *et al.*, 2019; Yassour *et al.*, 2016). Evaluating multiple sampling strategies—including the original data distribution, class weighting, and synthetic resampling—provides a more robust assessment of model reliability and reduces the risk of overfitting (Topçuoğlu *et al.*, 2020)

#### 2.4.7 Functional Profiling and Pathway Analysis

Functional metagenomic predictions were generated using PICRUSt2 (Douglas *et al.*, 2020), which infers gene family abundances from 16S rRNA data by leveraging ancestral-state reconstruction of known reference genomes. KEGG Orthologs (KOs) were predicted and mapped to MetaCyc and KEGG pathway hierarchies to enable downstream pathway-level analysis.

Differential pathway abundance between plant and animal protein diets was analyzed using MaAsLin2 (Mallick *et al.*, 2021), a multivariable linear modeling approach that allows covariate adjustment and correction for multiple testing. Relative abundances were log transformed, and protein source was included as a fixed effect. Pathways with false discovery rate (FDR)–adjusted p-values below 0.05 were considered statistically significant.

To visualize functional differences, heatmaps were generated using the pheatmap R package, focusing on the top-ranked differentially abundant pathways. Additional analysis focused on metabolic pathways related to sulfur metabolism, short chain fatty acid (SCFA) production, and nitrogen cycling. Taxa and pathway relationships were examined using Spearman correlation, and network topology metrics such as node centrality and clustering coefficients were used to characterize ecological interactions.

#### 2.4.8 Network Analysis

To investigate ecological interactions between microbial taxa and predicted functional pathways, correlation-based co-occurrence networks were constructed. Genus-level relative abundances and KEGG Orthologs predicted by PICRUSt2 were used as input. Pairwise Pearson correlation coefficients were computed, and significant associations were retained based on a p-value threshold of < 0.05 and an absolute correlation coefficient (|r|) ≥ 0.4, consistent with prior network-based microbiome studies (Kuntal *et al.*, 2019; Liu *et al.*, 2023).

Networks were visualized using the igraph and ggraph R packages. In each network, nodes represented microbial genera or functional pathways, and edges indicated significant positive or negative correlations. Edge thickness corresponded to the strength of the correlation.

To further characterize the structure of these networks, topological features were assessed using centrality measures, including *degree*, *betweenness*, and *eigenvector centrality*. Degree centrality reflects the number of direct connections a node has, indicating its immediate influence or connectivity within the network. Betweenness centrality captures how often a node lies on the shortest paths between other nodes, thus highlighting its role as a potential bridge or bottleneck in network flow. Eigenvector centrality assigns importance based not only on the number of connections but also on the influence of connected neighbors, identifying nodes with broader systemic influence. These measures were used to identify potential keystone taxa and central metabolic pathways that may play critical roles in shaping the microbiome’s structure and function (Faust & Raes, 2016; Liu *et al.*, 2019).

This integrative approach allowed for the simultaneous analysis of microbial community structure and functional potential, revealing clusters of co-associated taxa and pathways shaped by dietary protein source.

### 2.5 Cross-Species Comparison Between Human and Murine Microbiota

To evaluate whether microbial patterns identified in mice reflect broader trends in humans, we performed a cross-species comparison using stool microbiome data from the American Gut Project (AGP) (McDonald *et al.*, 2018), one of the largest publicly available human 16S rRNA datasets. Approximately 20,000 AGP samples were reprocessed using the same analytical workflow applied to the murine data. Human FASTQ files were denoised with DADA2 (Callahan *et al.*, 2016), taxonomic assignment was performed using the SILVA 138 reference database (Klindworth *et al.*, 2013), and centered log-ratio (CLR) transformation was applied to normalize compositional abundances (Gloor *et al.*, 2017; Quinn *et al.*, 2018). Matching these preprocessing steps minimized technical variability between species

Shared genera were identified from the CLR-transformed human and murine datasets and used to compute three complementary indicators of cross-species similarity: (1) mean relative abundance and prevalence of each shared genus in the AGP population, (2) correspondence in CLR-transformed abundance between species using Spearman correlation, and (3) genus-level effect-size differences expressed as ΔCLR (Human − Mouse). Murine CLR values were derived from the high-protein diet cohort to provide a diet-aligned reference for comparison. All analyses were performed in R (v4.4.1) using phyloseq (McMurdie & Holmes, 2012), microbiome (Lahti & Shetty, 2017), dplyr (Wen *et al.*, 2023), and ggplot2 (Wickham, 2016), with identical filtering and normalization steps applied across datasets to ensure methodological consistency.

## 3.0 Results

### 3.1 Characteristics of Selected Studies

The systematic search identified a total of 5,380 articles from PubMed (n = 339), EMBASE (n = 540), Scopus (n = 2,741), Web of Science (n = 1,710), and manual searches (n = 50). After de-duplication and preliminary screening, 2,696 abstracts were excluded due to duplicates (n = 982), review articles (n = 1,188), conference abstracts (n = 309), meta-analyses (n = 8), and letters or editorials (n = 209).

Subsequently, 2,684 full-text articles were assessed for eligibility. Of these, 2,668 were excluded for the following reasons: lack of an appropriate control group (n = 237), use of non-murine models (n = 1,761), inability to retrieve sequencing data (n = 534), or absence of a defined protein intervention (n = 136).

Ultimately, 10 studies met all inclusion criteria and were included in the meta-analysis (Masuoka *et al.*, 2020; Xie *et al.*, 2020; Ijaz *et al.*, 2020; Ruocco *et al.*, 2020; Choi *et al.*, 2021; Tan *et al.*, 2022; Yin *et al.*, 2022; Zhu *et al.*, 2022; Zhong *et al.*, 2022; Zhang *et al.*, 2022) **(Table 1).** These studies provided accessible 16S rRNA sequencing data, clearly defined animal or plant-based protein interventions, and complete sample metadata. To improve transparency in dietary comparisons, we summarized the macronutrient composition (%kcal from protein, fat, and carbohydrates including fiber) for all included rodent diets in **Supplementary Table 2**. Although nine distinct dietary protein sources were represented—branched chain amino acids (BCAAs), beef, casein, chicken, essential amino acids (EAAs), low protein diets, pork, sausage, and soy protein—they were categorized into two main groups for analysis: animal based proteins (beef, casein, chicken, pork, sausage, and BCAAs/EAAs when derived from animal sources) and plant based proteins (soy and low protein diets, as applicable). This binary classification enabled a focused comparison of microbial responses to protein origin across studies. A detailed breakdown of sample distribution by protein type and protein source is shown in **Supplementary Figure 1**. The study selection process is summarized in **Figure 1**.

**Table 2:** Core microbiota based on protein source at Genus level.

| Genera | Animal Protein (%) | Genera | Plant Protein (%) |
| --- | --- | --- | --- |
| *Muribaculaceae* | 21.93 | *Muribaculaceae* | 35.64 |
| *Faecalibaculum* | 10.01 | *Faecalibaculum* | 6.17 |
| *Akkermansia* | 8.70 | *Akkermansia* | 6.02 |
| *Lactococcus* | 7.34 | *Allobaculum* | 4.79 |
| *Ligilactobacillus* | 4.24 | *Dubosiella* | 4.79 |
| *Staphylococcus* | 3.25 | *Bacteroides* | 3.02 |
| *Bifidobacterium* | 3.21 | *Ligilactobacillus* | 2.99 |
| *Rikenellaceae_RC9_gut_* | 2.82 | *Alloprevotella* | 2.15 |

### 3.2 Composition of Core Gut Bacteria in Protein-fed mice

A total of 9,575,175 sequencing reads were generated from 187 protein fed mouse samples, yielding 16,630 amplicon sequence variants (ASVs), with an average of 51,204 sequence reads per sample. After denoising and chimera removal, 10,123 ASVs remained. Across all samples, Firmicutes (58.48%) and Bacteroidota (21.88%) were the dominant phyla, followed by Proteobacteria (7.08%), Actinobacteriota (4.13%), Acidobacteriota (1.34%), and Verrucomicrobiota (1.28%). The Firmicutes to Bacteroidota (F: B) ratio, a traditional indicator of metabolic health, was significantly higher in animal protein fed mice (63.3:19.9) compared to plant protein fed mice (47.3:23.5) (**Supplementary Table 3**, **Supplementary Figure 2**).

**Figure 2.**
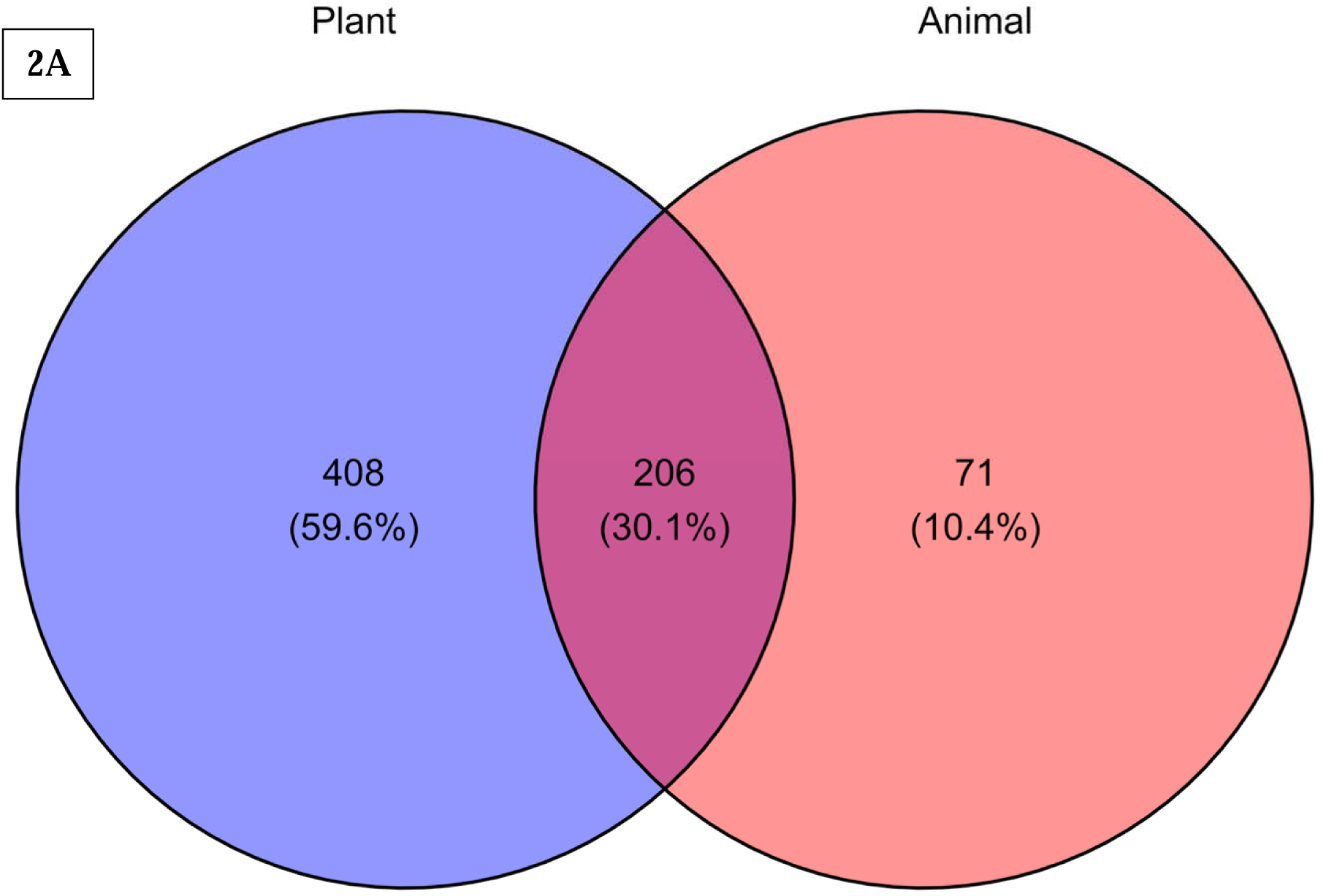
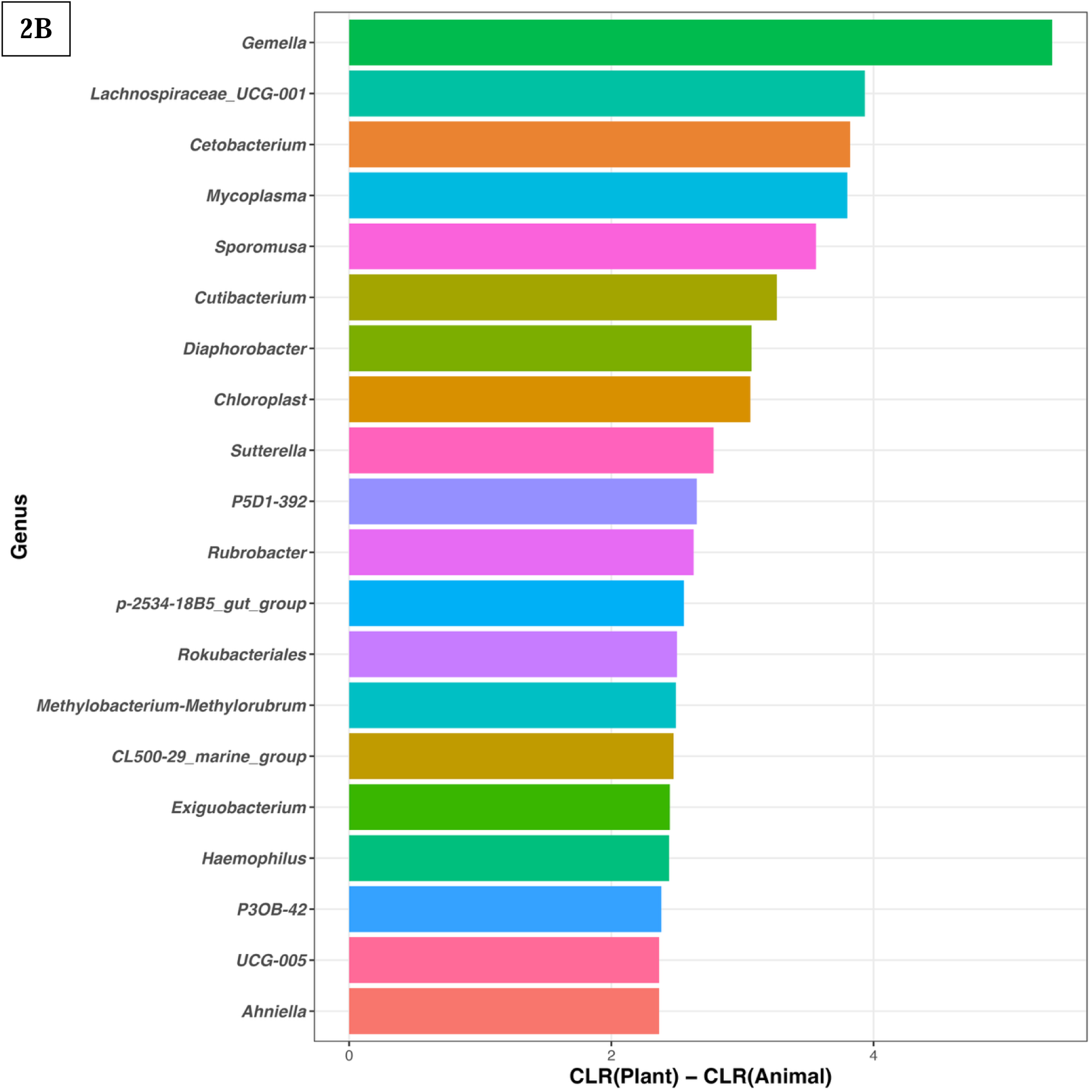
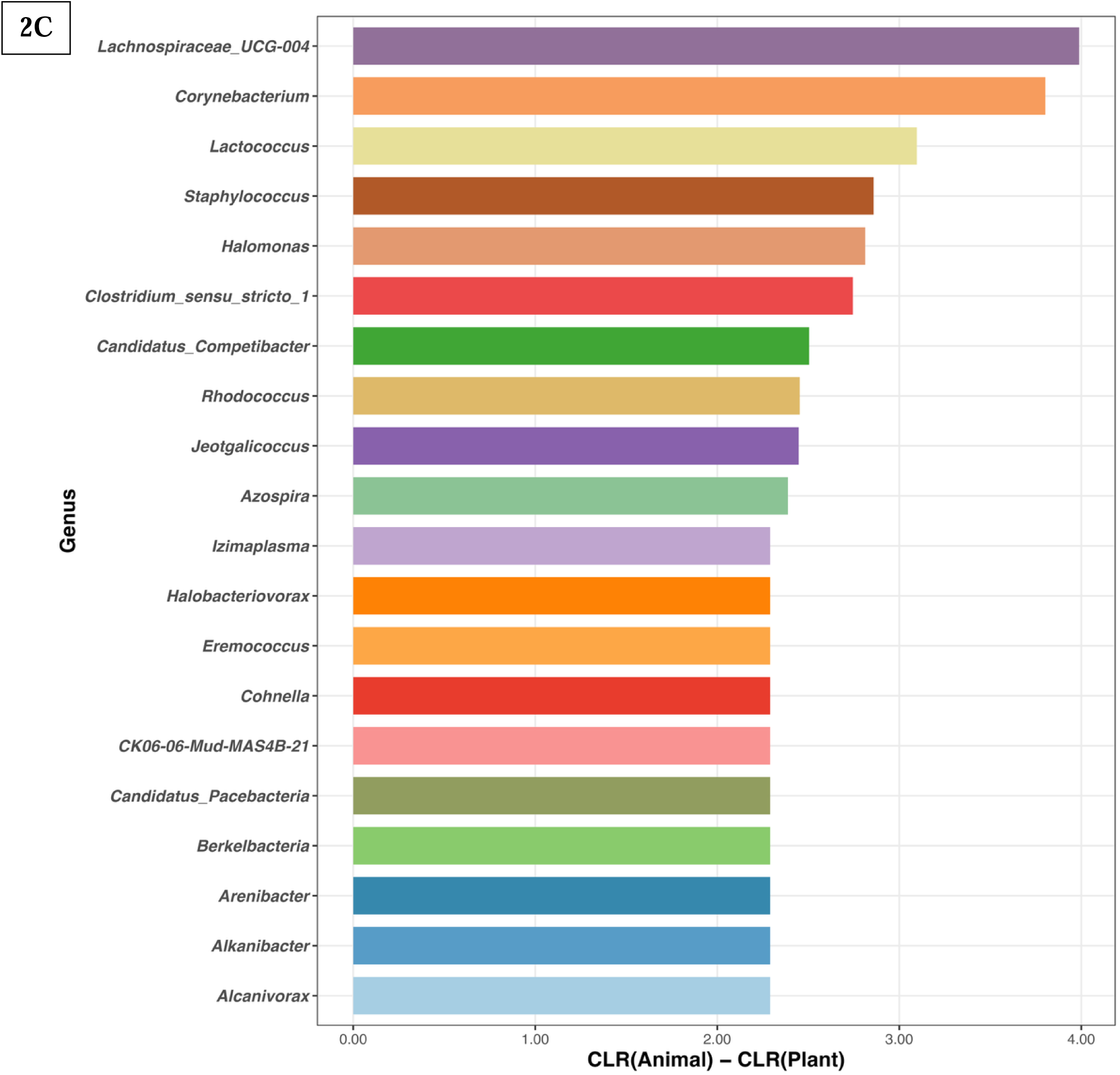
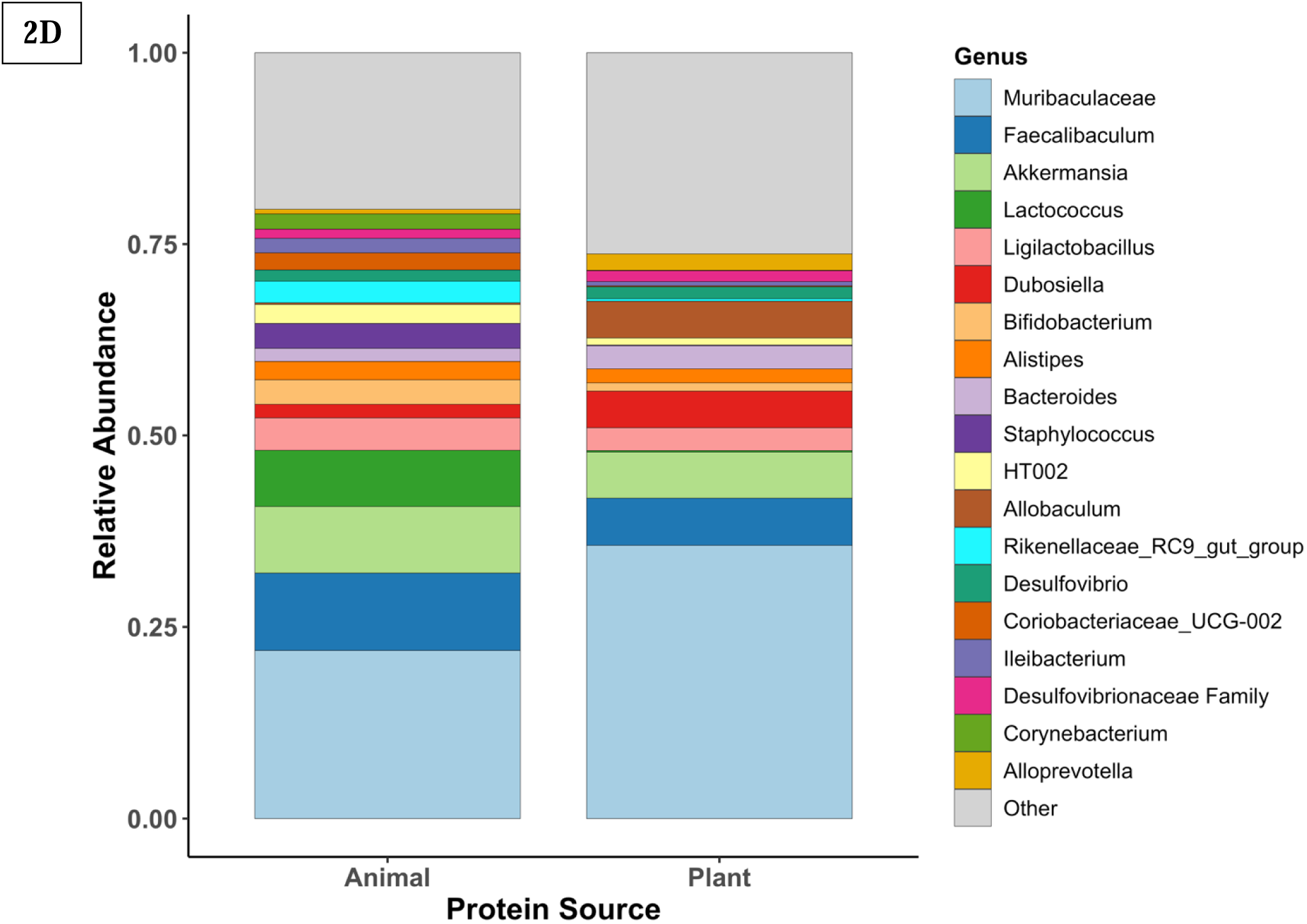
Microbial composition and taxonomic uniqueness across dietary protein sources. (A) Venn diagram showing the number and percentage of microbial genera unique to plant-fed and animal-fed mice, as well as genera shared between both groups. **(B)** Top genera enriched in plant-protein diets, displayed as CLR(Plant) − CLR(Animal) differences. Positive values indicate higher normalized abundance under plant protein intake **(C)** Top genera enriched in animal-protein diets, displayed as CLR(Animal) − CLR(Plant) differences. Positive values indicate higher normalized abundance under animal protein intake. **(D)** Stacked bar plot comparing the relative abundance of dominant genera across dietary groups, highlighting taxa that differ consistently between plant- and animal-protein diets

Among the 685 identified genera, plant protein diets supported a significantly higher number of unique microbial taxa (408 genera, 59.6%) compared to animal protein diets (71 genera, 10.4%) (**Figure 2A**), indicating greater microbial diversity. Despite this difference, both dietary groups shared a core set of dominant taxa, with *Muribaculaceae*, *Faecalibaculum*, and *Akkermansia* highly abundant across all samples, although their relative abundance varied by diet.

To quantify how unique taxa were distributed across diets, we evaluated centered log-ratio (CLR) enrichment among genera detected exclusively within each group. Plant-unique taxa exhibited uniformly positive CLR differences (CLR*^Plant^* − CLR*^Animal^*), indicating consistent enrichment under plant-protein feeding **(Figure 2B).** The most strongly represented genera included *Gemella, Lachnospiraceae UCG-001, Cetobacterium, Mycoplasma,* and *Sporomusa,* along with several low-abundance fermentative taxa that contributed to the broader ecological repertoire of plant-fed microbiota. Conversely, genera unique to animal-protein diets displayed positive CLR differences in the opposite direction (CLR*^Animal^* − CLR*^Plant^*) reflecting preferential representation under animal-derived proteins **(Figure 2C).** Key contributors included *Lachnospiraceae UCG-004, Corynebacterium, Lactococcus, Staphylococcus, Halomonas,* and *Clostridium sensu stricto 1,* many of which are associated with proteolytic metabolism.

Beyond unique genera, relative abundance comparisons of shared taxa revealed clear dietary signatures **(Figure 2D**, **Table 2).** Plant-protein diets were characterized by higher levels of saccharolytic fermenters, including *Muribaculaceae* (35.64% vs. 21.93% in animal-fed mice), *Dubosiella* (4.79% vs. 2.11%), and *Allobaculum* (4.79% vs. 1.42%), reflecting enhanced utilization of plant-derived substrates. In contrast, animal-protein diets showed increased representation of proteolytic taxa such as *Lactococcus* (7.34% vs. 0.82% in plant-fed mice), *Ligilactobacillus* (4.24% vs. 1.03%), and *Staphylococcus* (3.25% vs. 0.56%), genera commonly associated with protein fermentation and nitrogen-rich metabolic byproducts. These differences highlight distinct compositional patterns associated with each protein source, with plant-derived proteins showing higher abundance of saccharolytic taxa and animal-derived proteins showing greater representation of proteolytic genera.

### 3.3 Dietary Protein Sources Modulate Microbial Diversity

To assess how dietary protein source, protein concentration, and other contextual factors affect microbial diversity, we employed a generalized linear mixed model (GLMM) framework as described in Section 2.4.2. The best-fitting model included dietary protein source and protein concentration as fixed effects, with random intercepts specified for country of origin, sample isolation site, and animal age (in weeks). This model achieved the lowest AIC (252.88) and BIC (278.20), and the highest log-likelihood (−118.44) **(Supplementary Table 4).** Model selection was further validated by likelihood ratio tests and R² metrics, revealing a marginal R² of 0.071 and a conditional R² of 0.893, indicating substantial contribution from both fixed and random effects **(Figure 3A).** Protein source had the strongest positive association with Shannon diversity (β = 1.25), followed by protein level (β = 0.40). Sequencing depth did not significantly impact diversity estimates (ρ = 0.107, p = 0.145), suggesting biological rather than technical drivers of variation **(Supplementary Figure 3A).**

**Figure 3:**
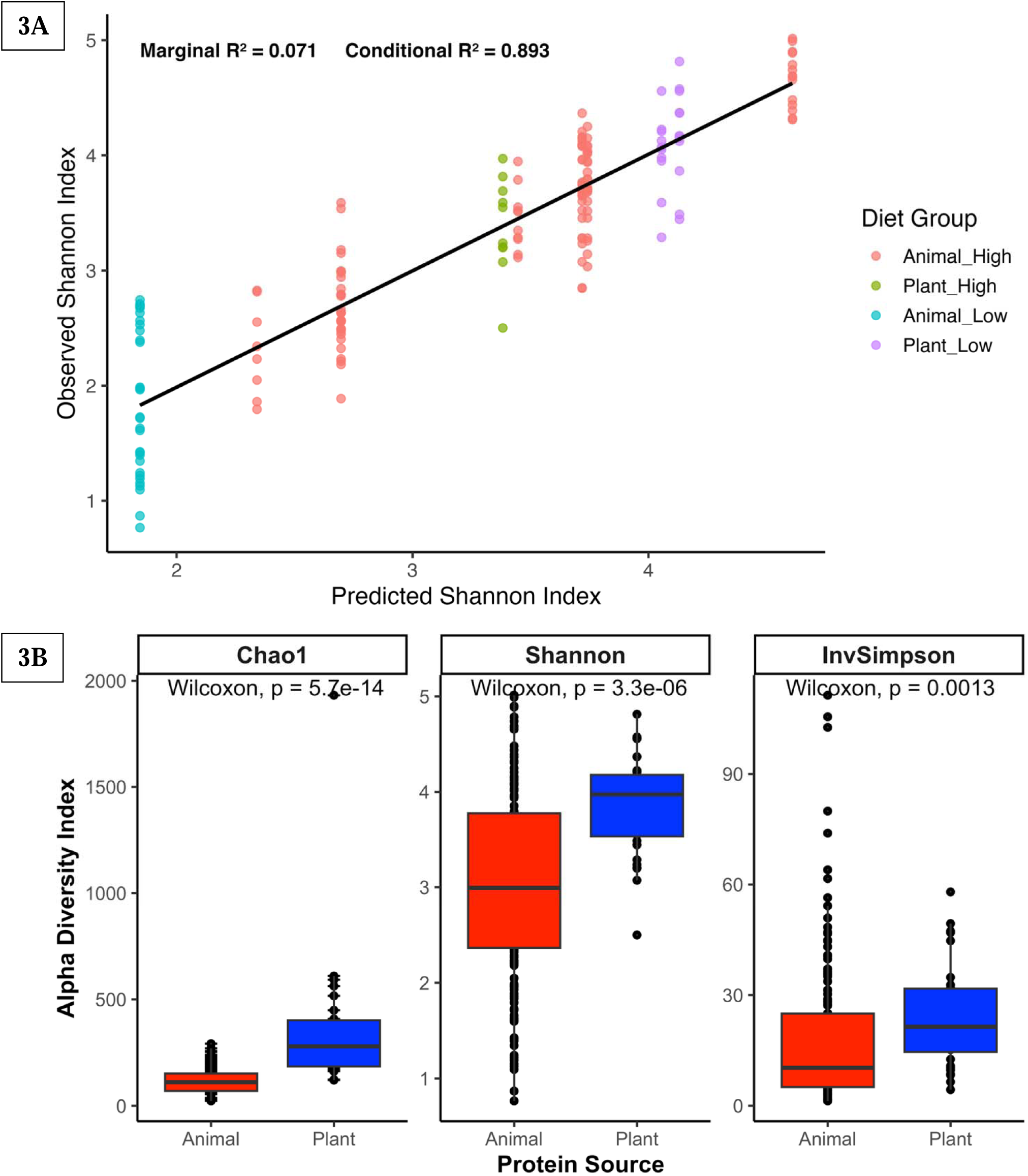
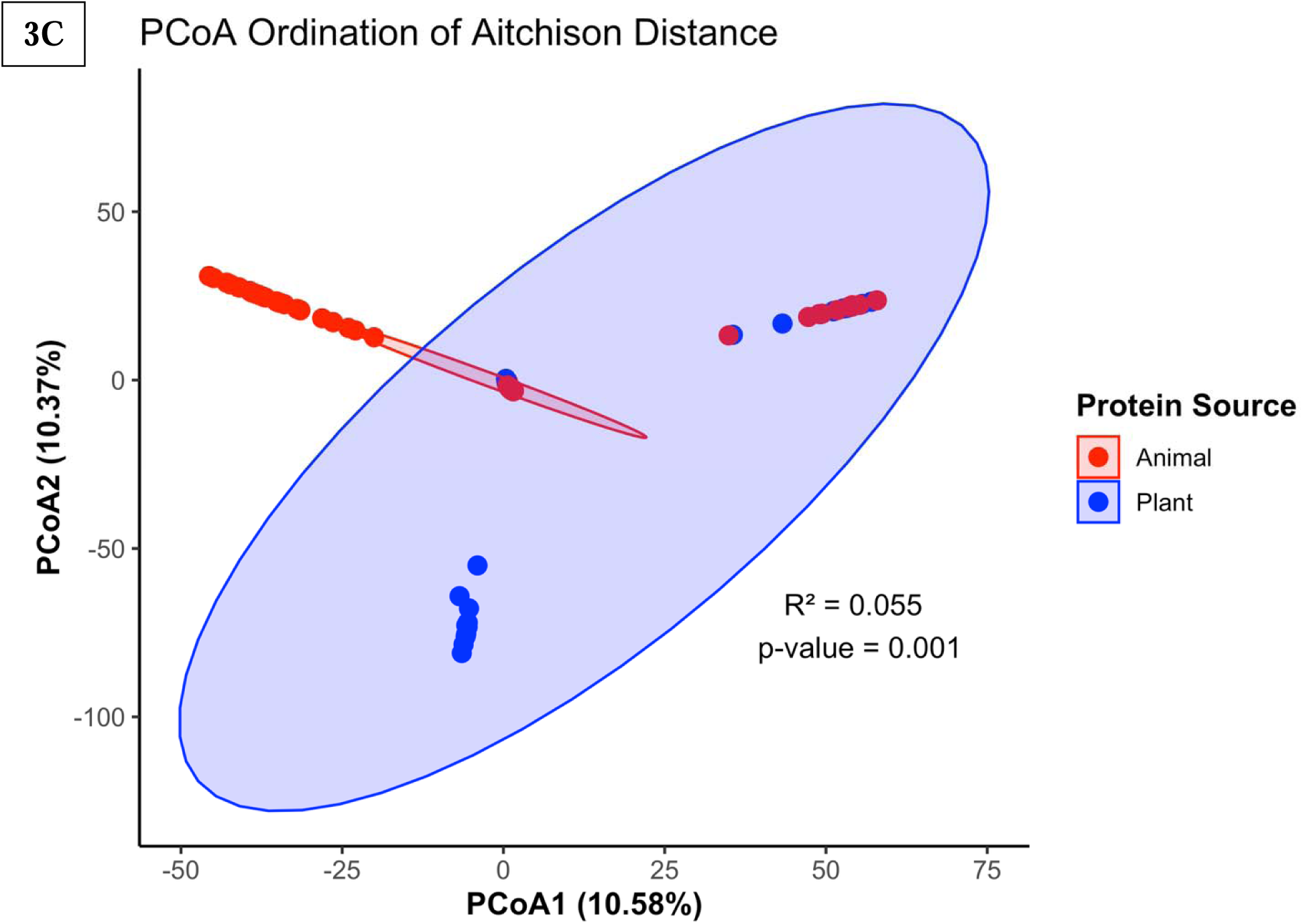
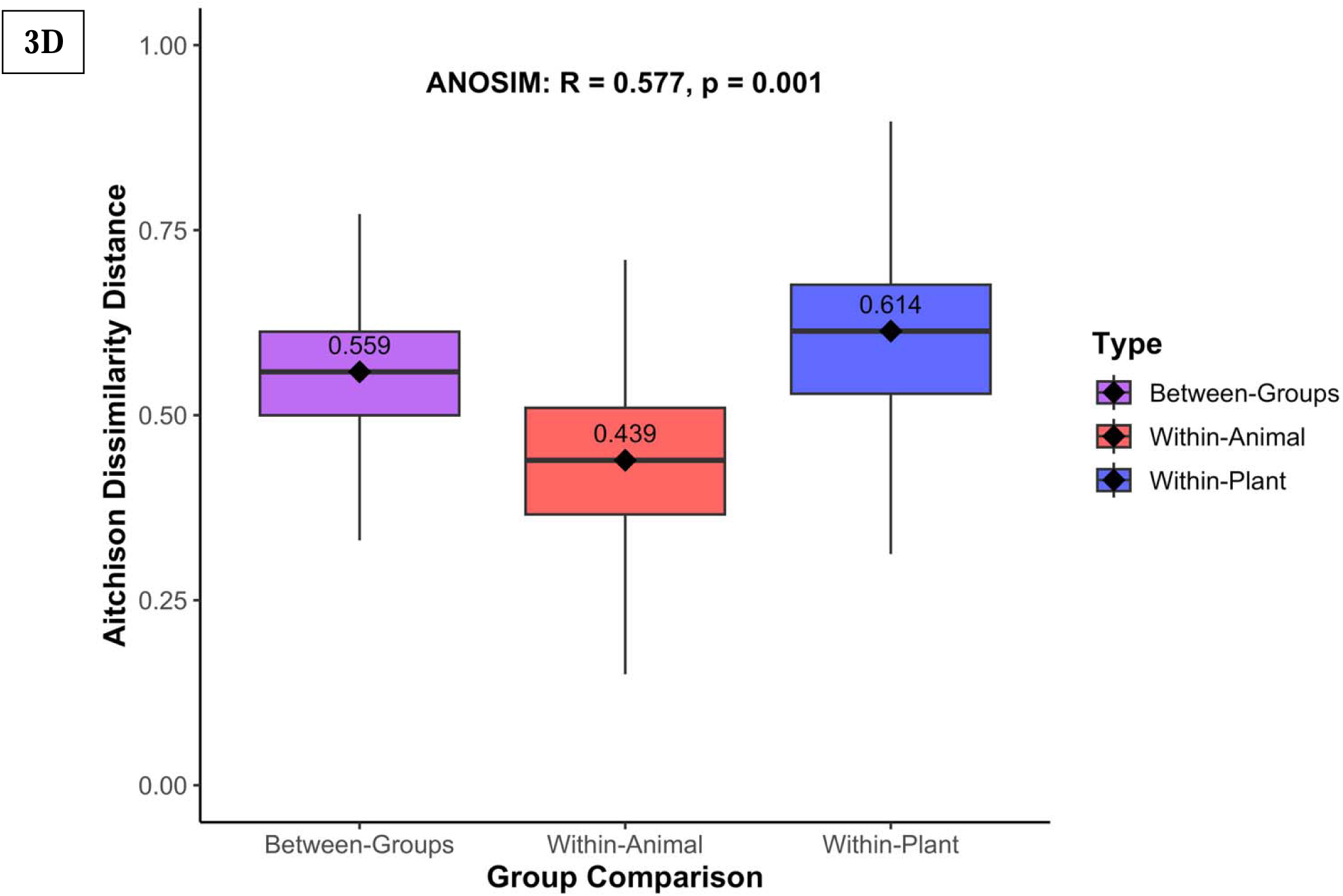
Microbial Diversity and Community Structure in Protein-Based Diets. (A) Linear mixed model comparing predicted versus observed Shannon diversity, with marginal and conditional R² values indicating variance explained by fixed effects alone (0.071) versus fixed plus random effects (0.893), respectively (B) Boxplots showing significantly higher microbial richness and evenness in plant-based protein diets. (C) Principal Coordinates Analysis (PCoA) based on Aitchison distance reveals clear separation of microbial communities by protein source. Ellipses represent 95% confidence intervals. (D) ANOSIM analysis confirms significant differences in microbial community structure between diet groups (R = 0.663, P < 0.001).

Consistent with these model outputs, microbial alpha diversity was significantly higher in plant protein diets across all three diversity indices: Chao1 (*p* = 5.7 × 10⁻¹⁴), Shannon (*p* = 3.3 × 10⁻⁶), and Inverse Simpson (*p* = 0.0013) (**Figure 3B**). A notable interaction between protein source and concentration was detected **(Supplementary Figure 3B),** with Shannon diversity highest in high plant-protein diets (*p* = 7.39 × 10⁻¹⁵), while differences were not significant at low protein levels (*p* = 0.722). These results suggest that both the source and concentration of dietary protein contribute to microbial richness and evenness.

To further examine shifts in overall microbial community composition, we analyzed beta diversity using Aitchison distance-based Principal Component Analysis (PCA). The ordination revealed distinct clustering by protein source, with the first two components explaining 10.58% and 10.37% of the total variance, respectively (**Figure 3C**). These compositional differences were confirmed by PERMANOVA (*R²* = 0.055, *p* = 0.001) and ANOSIM (*R* = 0.663, *p* = 0.001), underscoring significant divergence in microbial communities between plant- and animal-protein diets (**Figure 3D**).

Together, these findings demonstrate that dietary protein source exerts a dominant influence on both within-sample (alpha) and between-sample (beta) microbial diversity, independent of sequencing depth or study-specific factors. High plant protein intake supports a more diverse and compositionally distinct gut microbiota, implicating protein source as key in modulating the gut microbial ecosystem.

### 3.4 Machine Learning Identifies Microbial Predictors of Protein Source

Supervised machine learning confirmed that gut microbial composition is highly predictive of dietary protein source. Among the three Random Forest (RF) models evaluated, the unadjusted RF yielded the highest classification performance (AUC = 1.00, Accuracy = 98%), followed by the ROSE-sampled RF (AUC = 0.997, Accuracy = 92.2%) and the class-weighted RF (AUC = 0.995, Accuracy = 88.2%) (**Figure 4A**).

**Figure 4.**
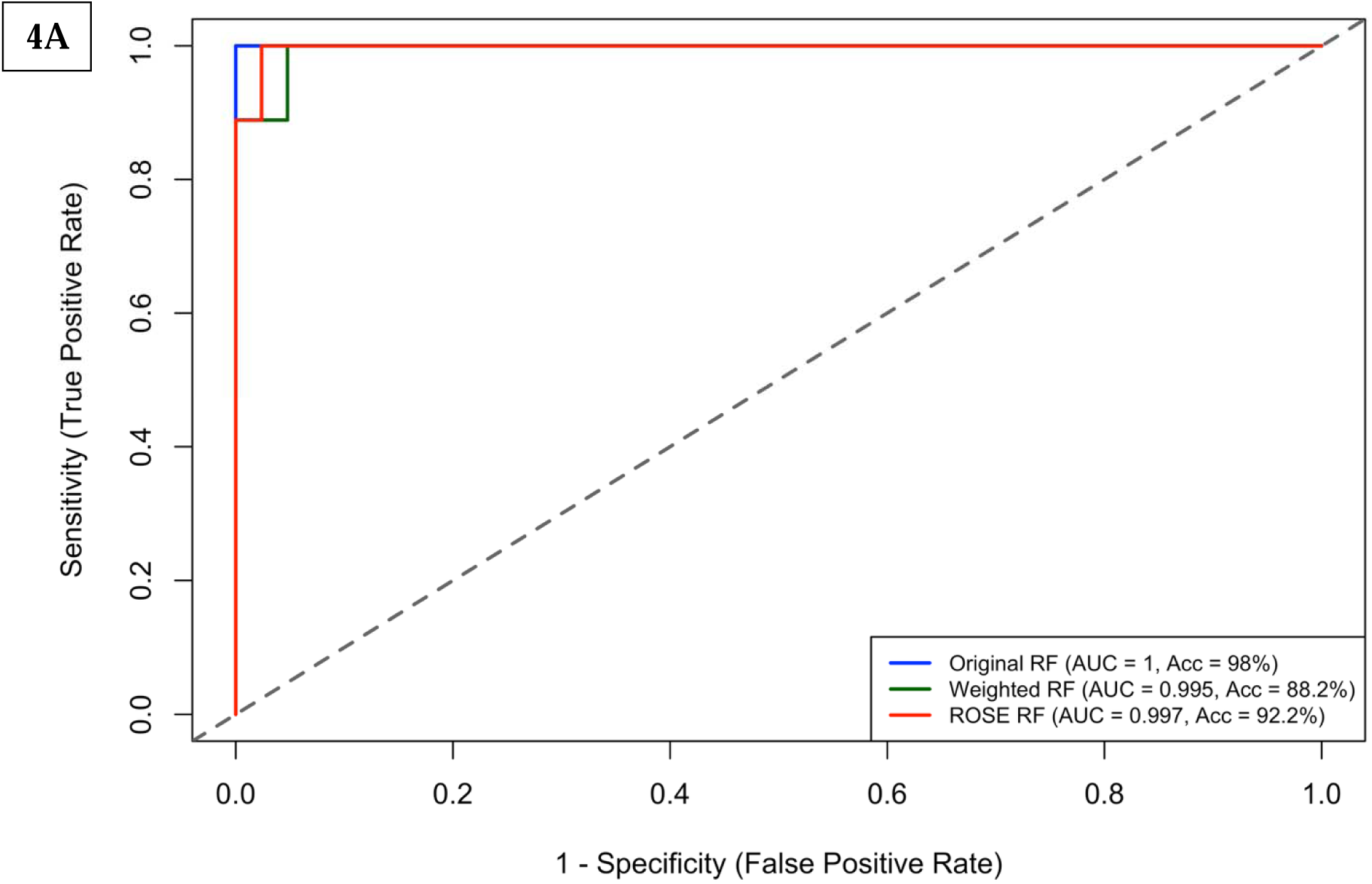
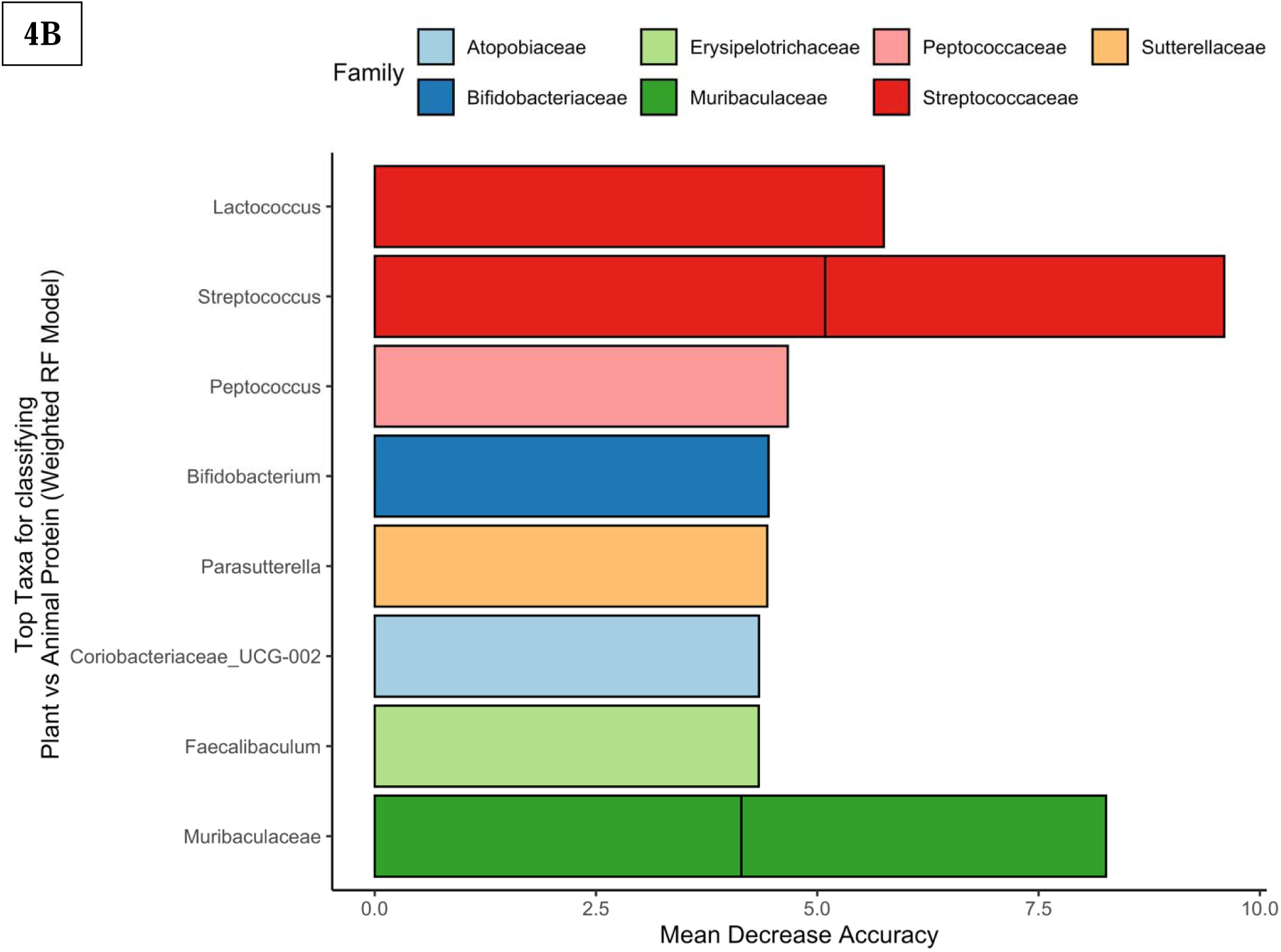
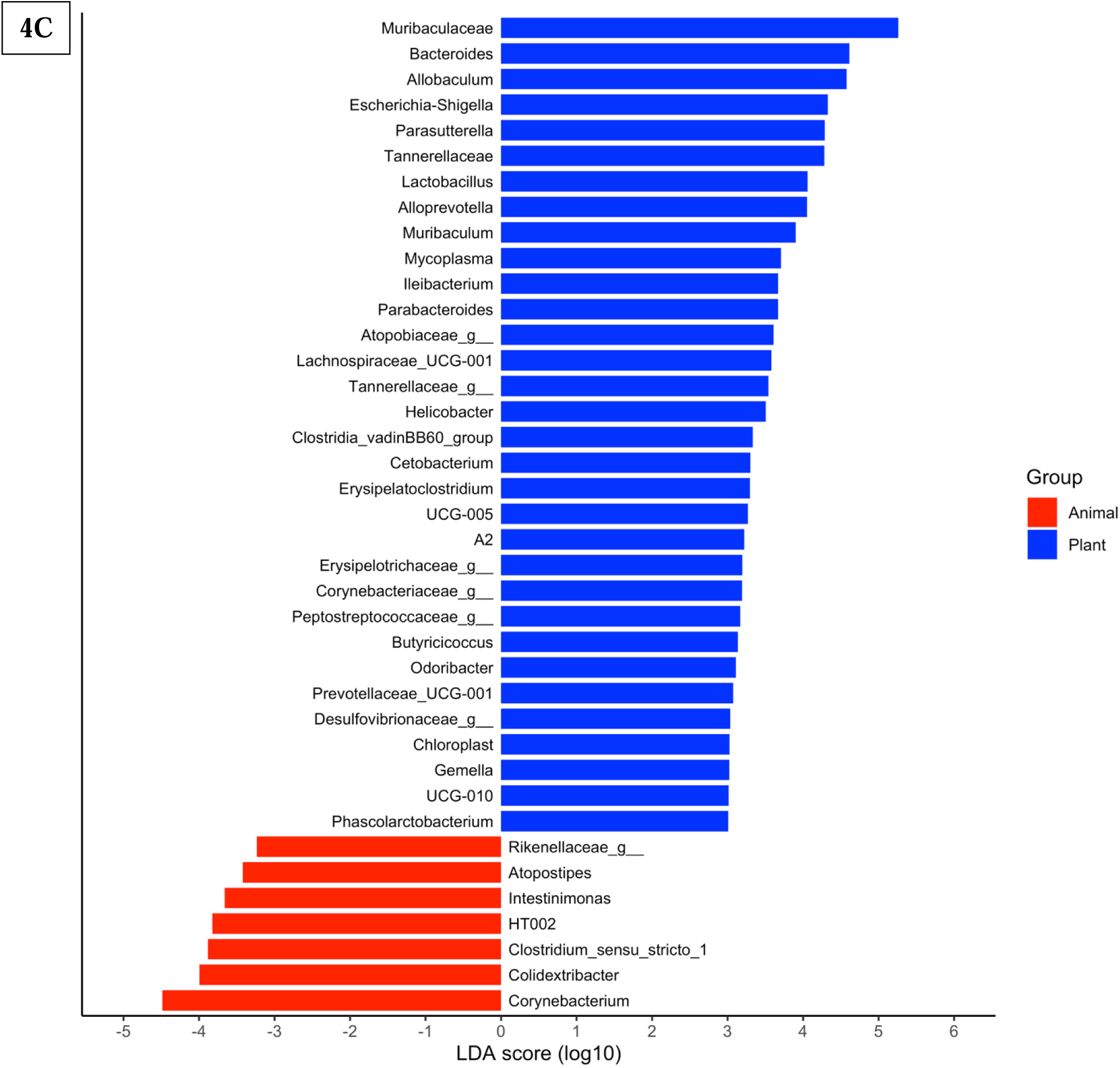
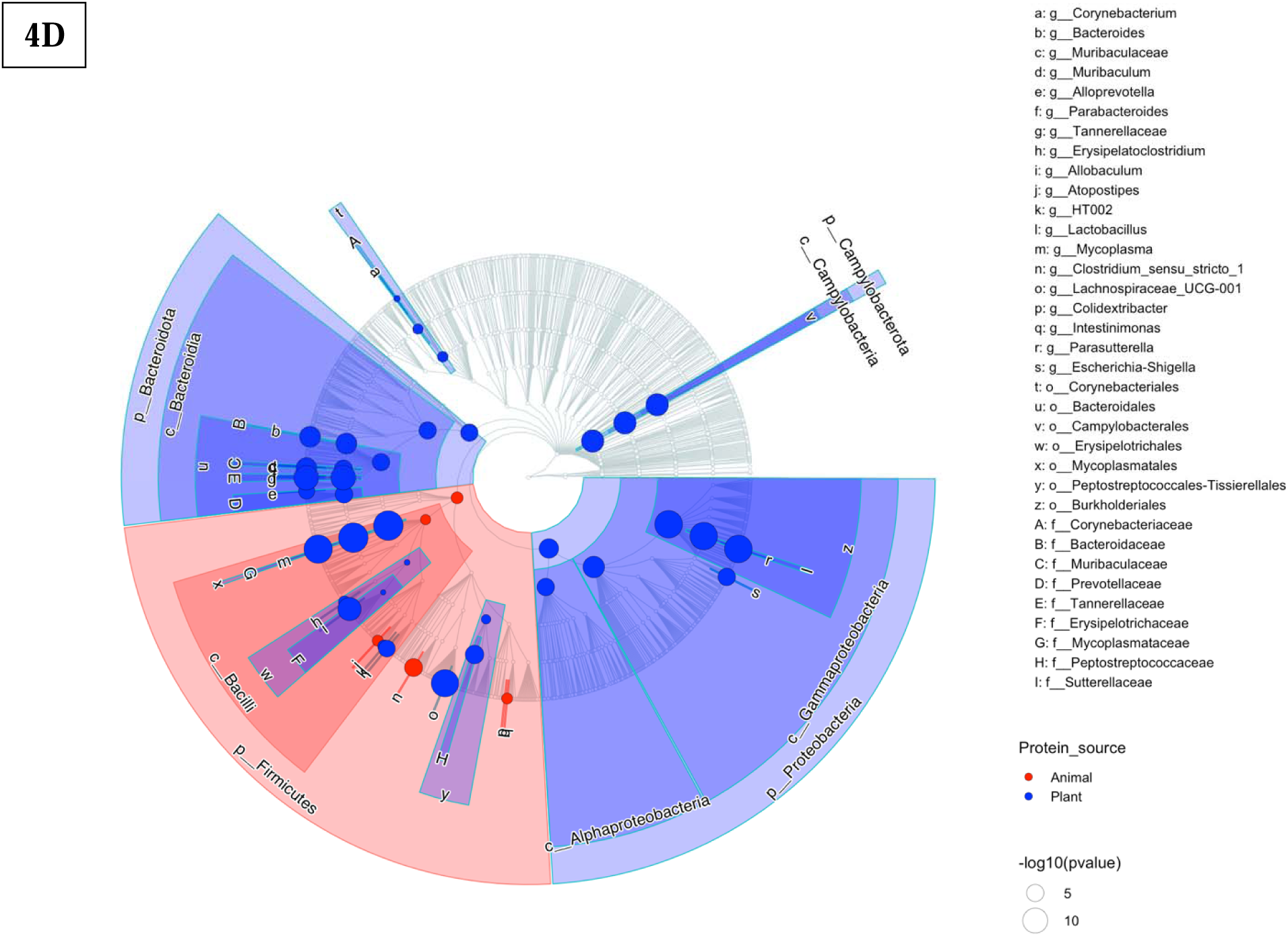
Machine Learning performance and key microbial predictors of dietary protein source. (A) Receiver Operating Characteristic (ROC) curves showing percentage accuracy and area under curve (AUC) comparing three Random Forest (RF) models: unadjusted (Original), class-weighted (Weighted), and balanced via ROSE sampling. The weighted RF model, while slightly lower in accuracy, avoids overfitting and offers more realistic performance, with an AUC of 0.995 and accuracy of 88.2%. (B) Top 10 microbial genera contributing most to the weighted RF model’s classification of animal vs plant protein diets, based on Mean Decrease in Accuracy (MDA). (C). LDA using LEfSe highlights taxa enriched in each diet (LDA score ≥ 3.0, p < 0.05) (D). Cladogram shows microbial taxa enriched in plant-based (blue) and animal-based (red) diets, arranged hierarchically from phylum to genus with node sizes reflecting statistical significance (-log10[p-value]).

Despite its slightly lower accuracy, the class-weighted model was selected for downstream feature selection to correct for class imbalance between plant- and animal-protein groups. Feature importance analysis based on Mean Decrease Accuracy (MDA) identified key taxa contributing to classification, including *Lactococcus* (5.34%), *Streptococcus* (5.09%, 4.34%), *Peptococcus* (4.71%), *Bifidobacterium* (4.56%), *Parasutterella* (4.47%), *Faecalibaculum* (4.43%), *Coriobacteriaceae UCG-002* (4.44%), and *Muribaculaceae* (4.10%, 4.06%) **(Figure 4B; Supplementary Table 5).**

Model robustness was supported by cross-validation and comparison with a randomized control model, which achieved only 38.2% accuracy **(Supplementary Figure 4A).** The error-per-tree diagnostic further confirmed model stability across 1,000 trees **(Supplementary Figure 4B).** Confusion matrix analysis showed high specificity for plant-protein predictions, with modest misclassification among animal-protein samples **(Supplementary Figure 4C).**

LEfSe analysis further validated these findings, identifying 33 genera enriched in plant-protein diets and 7 in animal-protein diets. Notably, plant-associated genera included *Muribaculaceae, Bacteroides, Parabacteroides,* and *Allobaculum*, while animal-protein diets were characterized by genera such as *Clostridium sensu stricto* 1*, Colidextribacter,* and *Rikenellaceae.* A LEfSe-derived cladogram illustrated the phylogenetic structure of these associations, with plant-protein diets favoring Bacteroidota and Proteobacteria, and animal-protein diets skewed toward Firmicutes dominance **(Figure 4C–D; Supplementary Table 6).** These results demonstrate the use of machine learning for uncovering diet-responsive microbial biomarkers and highlight the biological relevance of microbial shifts linked to dietary protein type.

### 3.5 Functional Pathways Enriched by Protein Type

Functional profiling using PICRUSt2 revealed distinct metabolic signatures between plant- and animal-protein diets. Differential pathway enrichment analysis identified 10 plant-enriched and 10 animal-enriched MetaCyc pathways with significant log₂ fold changes **(Figure 5A).** Plant-based diets were associated with pathways related to carbohydrate metabolism and amino acid degradation, including the Entner-Doudoroff pathway, nitrate reduction, aromatic biogenic amine degradation, and L-tryptophan degradation. Conversely, animal-protein diets were enriched in fermentation and biosynthesis pathways such as creatinine degradation, propanoate fermentation, and L-isoleucine biosynthesis. These findings highlight distinct metabolic capacities of the microbiota shaped by dietary protein source.

**Figure 5.**
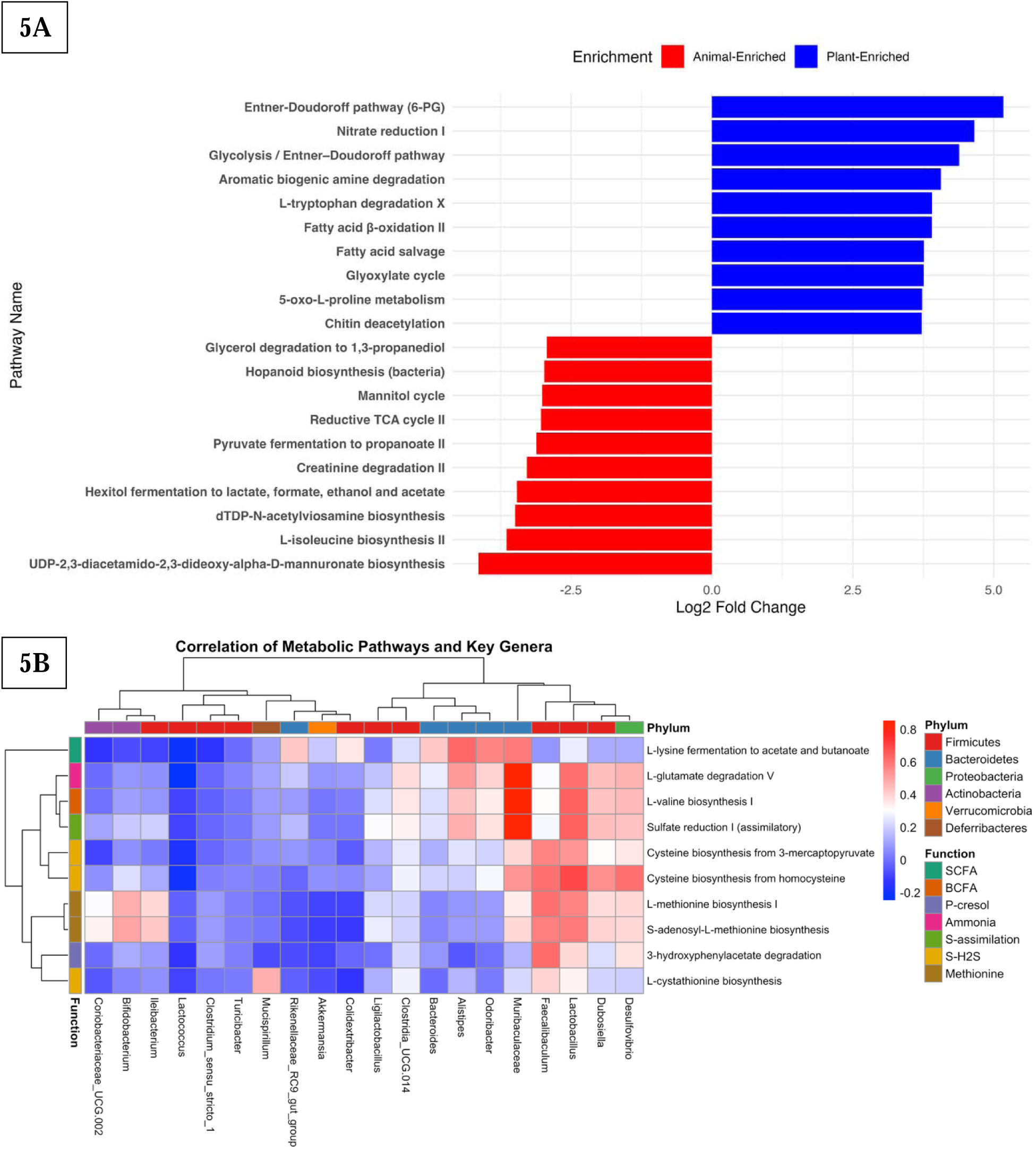
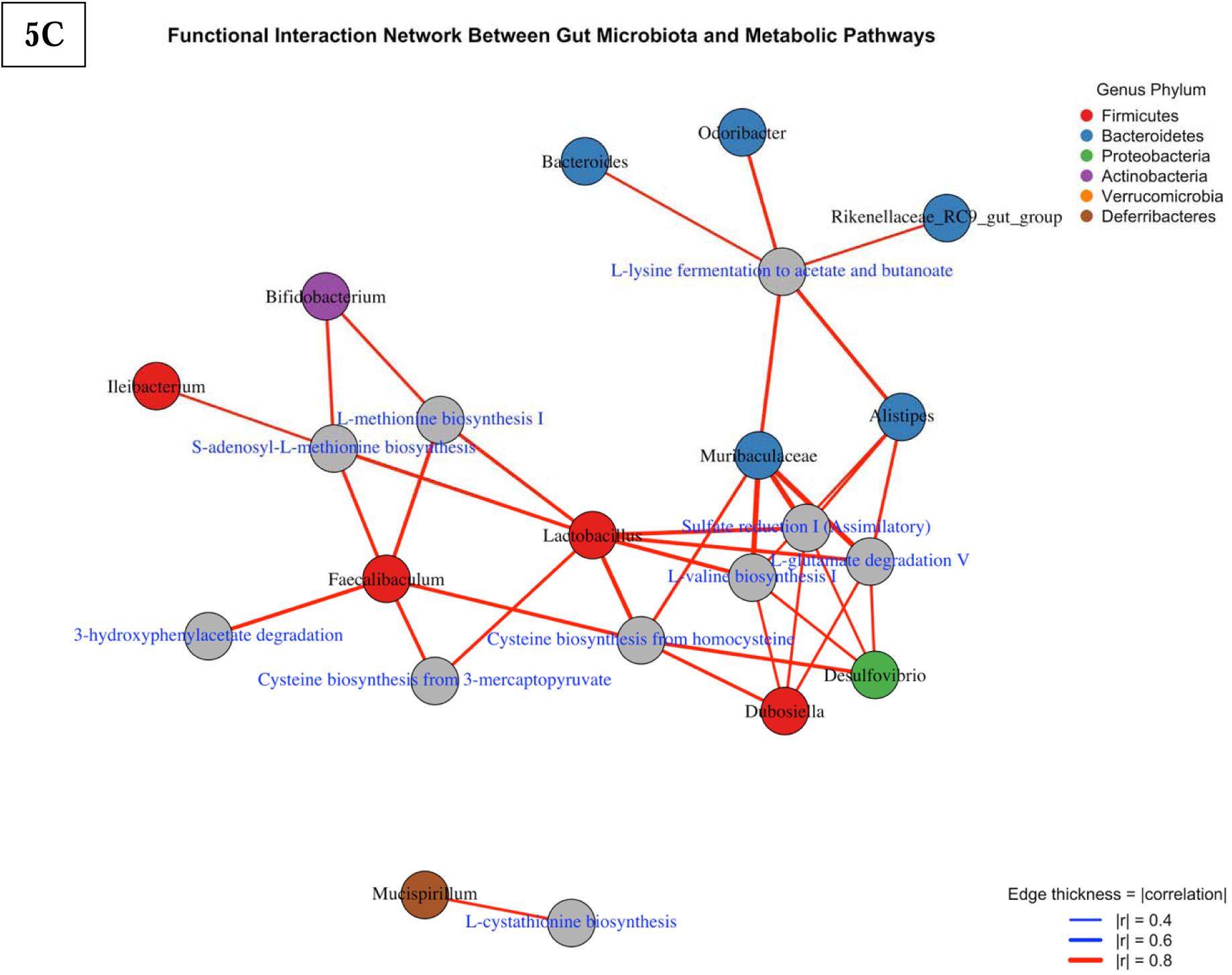
Functional profiling of microbial communities by protein source using PICRUSt2 predictions. (A) Top 10 KEGG pathways significantly enriched in mice fed animal- or plant-based protein diets (|log₂ fold change| > 2, FDR < 0.05). Red bars indicate enrichment in the animal group; blue bars indicate enrichment in the plant group. (B) Heatmap showing Pearson correlation between microbial genera and selected predicted metabolic pathways involved in short-chain fatty acid (SCFA), branched-chain fatty acid (BCFA), phenol/p-cresol, ammonia, and sulfur metabolism. Functions are color-coded by metabolic category, and genera are annotated by phylum. (C) Network diagram of strong correlations (|r| ≥ 0.6) between microbial genera and selected predicted metabolic pathways. Node size reflects the number of connections (degree centrality), and edge thickness corresponds to the strength of correlation.

### 3.6 Correlation and Network Analysis Reveal Functional Interactions

To investigate how enriched pathways relate to specific microbial taxa, a correlation analysis was performed between predicted KEGG Ortholog functions and the top differential genera. The resulting heatmap **(Figure 5B)** highlighted strong associations between specific microbial genera and functional pathways involved in SCFA, BCFA, sulfur, and amino acid metabolism. *Lactobacillus* correlated with cysteine biosynthesis (r = 0.693, *p* = 6.5×10⁻²⁸), sulfate reduction (r = 0.647, *p* = 7.1×10⁻²³), and methionine biosynthesis (r = 0.566, *p* = 2.4×10⁻¹⁷). *Faecalibaculum* showed positive correlations with methionine biosynthesis (r = 0.606, *p* = 1.4×10⁻²⁰) and cysteine biosynthesis (r = 0.602, *p* = 2.7×10⁻²⁰). *Dubosiella* was linked to glutamate degradation (r = 0.440, *p* = 2.5×10⁻¹⁰) and cysteine biosynthesis (r = 0.537, *p* = 1.8×10⁻¹⁵). *Bifidobacterium* was moderately correlated with methionine pathways, including S-adenosylmethionine biosynthesis (r = 0.503, *p* = 3.8×10⁻¹³). *Alistipes* and *Muribaculaceae* were associated with SCFA-related pathways, including lysine fermentation (*r* = 0.619, *p* = 8.5×10⁻²² and *r* = 0.582, *p* = 5.6×10⁻¹⁹, respectively), glutamate degradation (*r* = 0.509, *p* = 2.3×10⁻¹⁴ and *r* = 0.855, *p* = 2.1×10⁻⁵⁴), and valine biosynthesis (*r* = 0.422, *p* = 4.3×10⁻¹⁰ and *r* = 0.858, *p* = 7.6×10⁻⁵⁵). These patterns suggest that distinct microbial taxa specialize in different metabolic niches depending on dietary protein source.

Network analysis based on pairwise Pearson correlations (|r| ≥ 0.4, p < 0.05) visualized the co-occurrence structure of microbial taxa and metabolic functions **(Figure 5C).** Nodes represented taxa or pathways, and edge thickness indicated correlation strength. Highly connected nodes such as *Faecalibaculum* (eigenvector centrality = 0.97) and *Lactobacillus* (0.97) exhibited strong topological influence within sulfur and methionine metabolic subnetworks, suggesting they may serve as functional keystone taxa in modulating amino acid–related metabolic functions.

Similarly, *Muribaculaceae* demonstrated the highest eigenvector centrality (1.00) **(Supplementary Figure 5),** underscoring its potential integrative role across diverse metabolic modules, particularly those involved in glutamate degradation and SCFA production. These network metrics reinforce the centrality of select taxa in shaping key nutrient pathways influenced by dietary protein source.

### 3.7 Cross-Species Comparison of Shared Gut Bacterial Genera

A subset of genera detected in the murine datasets also appeared in the AGP cohort, demonstrating that several protein-responsive taxa are not restricted to mice. Among shared genera, *Brevundimonas* (24.6% prevalence) and *Tuzzerella* (7.8%) were the most frequently observed in humans, with additional contributions from *Weissella, Sphingomonas, Sphingopyxis, Christensenella,* and *Eubacterium* **(Figure 6A).**

**Figure 6.**
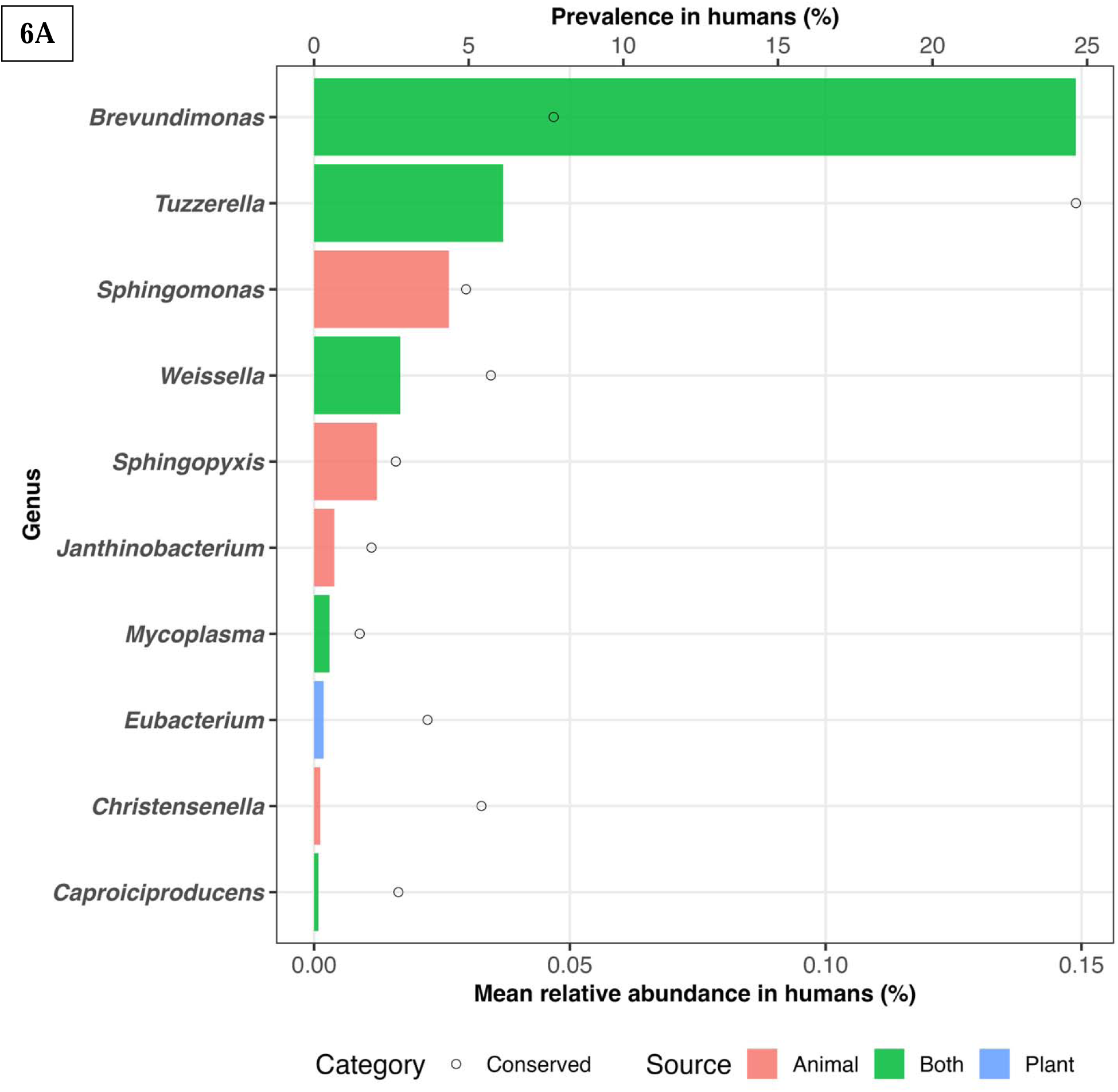
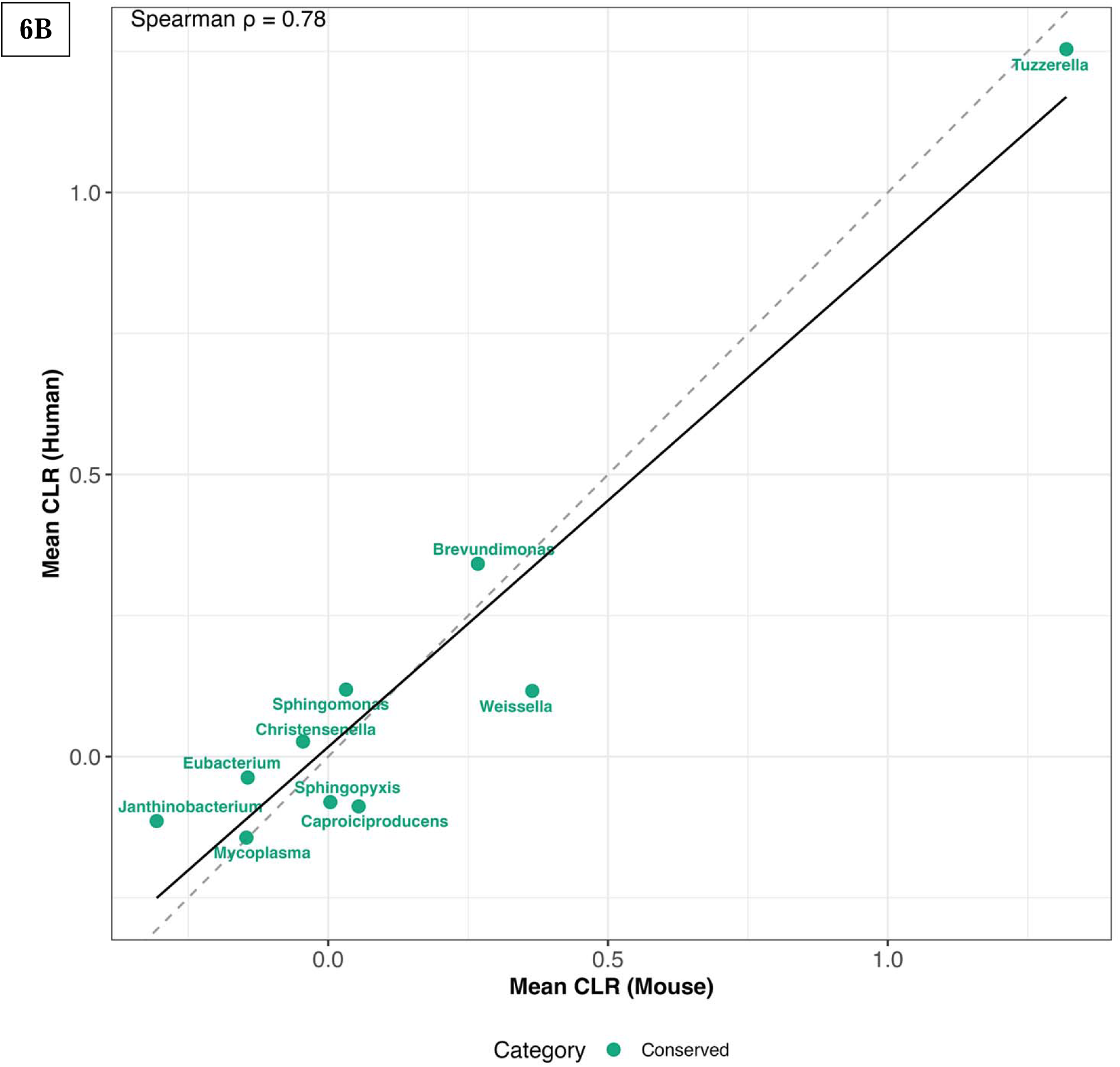
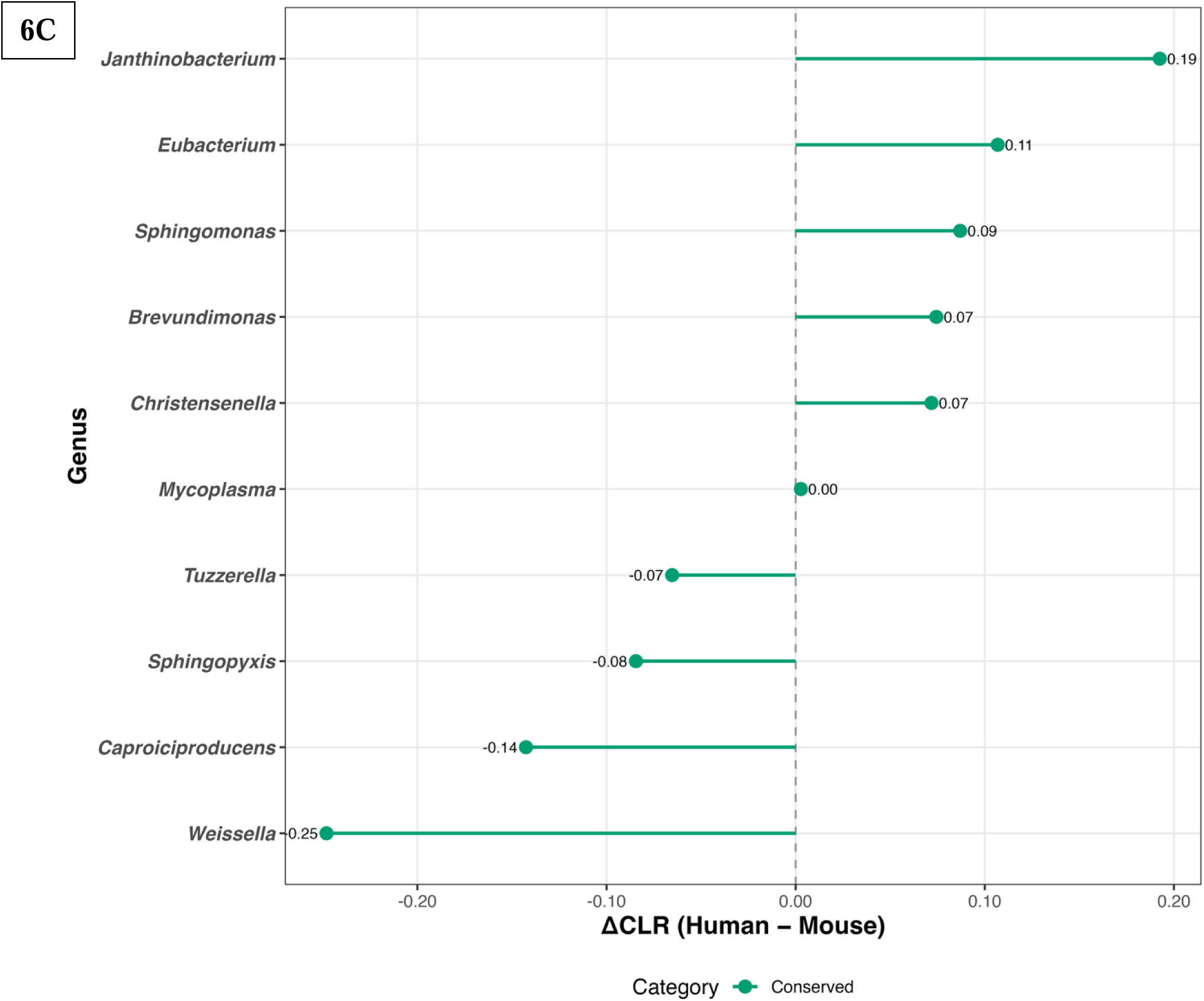
Cross-species compositional similarity of gut bacterial genera shared between humans and mice. **(A)** Mean relative abundance and prevalence of genera detected in both humans and mice; bars represent human relative abundance grouped by source type (animal-, plant-associated, or mixed), and points indicate human population prevalence. **(B)** Cross-species mean CLR per genus; points represent shared genera in humans and mice. The dashed diagonal denotes perfect correspondence (y = x), and the solid line shows the regression trend (Spearman ρ = 0.78). **(C)** CLR effect-size differences (ΔCLR = Human − Mouse). The lollipop plot displays direction and magnitude of cross-species shifts, where values near zero indicate similar ecological representation in both hosts.

CLR-transformed abundance profiles showed a positive cross-species correspondence (Spearman ρ = 0.78), indicating that genera with higher CLR values in mice tended to occupy similar positions along the abundance gradient in humans **(Figure 6B).**

Genus-level effect-size differences (ΔCLR = Human − Mouse) were modest and generally centered near zero **(Figure 6C).** *Janthinobacterium* and *Eubacterium* showed slightly elevated CLR values in humans, whereas *Weissella, Caproiciproducens,* and *Sphingopyxis* displayed higher CLR values in mice. Despite these differences, most genera fell within overlapping CLR ranges, reflecting broad consistency in their comparative abundance across hosts.

Together, these results indicate that several taxa enriched under protein-modulated conditions in mice are also represented in human stool samples, with broadly similar abundance patterns across species.

## 4.0 Discussion

This meta-analysis integrates 16S rRNA datasets across multiple murine studies to provide a comprehensive overview of how dietary protein sources are associated with gut microbial composition, diversity, and functional potential. In addition to compositional differences, we identify consistent ecological and metabolic patterns linked to saccharolytic versus proteolytic fermentation, highlighting microbial taxa and pathways that co-vary with protein source (De Filippis *et al.*, 2016; Canfora *et al.*, 2019). These findings deepen our understanding of how variation in dietary nitrogen and carbon availability corresponds with shifts in microbial community structure and function, with potential implications for host health.

### 4.1 Overview of Diet-Associated Taxonomic Patterns

Dietary protein source was strongly associated with distinct taxonomic profiles in the gut microbiota. Plant-derived proteins were associated with a broader range of taxa, including genera with saccharolytic and low-nitrogen metabolic capacities, consistent with the wider ecological niches supported by plant-feeding. In contrast, animal-derived proteins favored genera associated with proteolytic metabolism and nitrogen-rich substrates

CLR-based enrichment patterns further highlighted this divergence, showing that plant-associated genera generally exhibited higher CLR abundance under plant-protein feeding, whereas animal-associated genera showed the reverse pattern. These patterns are consistent with established ecological models describing how dietary nitrogen availability shapes competitive interactions and niche partitioning in the gut microbiome (Windey *et al.*, 2012; Neis *et al.*, 2015).

Overall, these taxonomic differences provide an ecological foundation for the subsequent diversity, biomarker, and functional pathway results, indicating that protein source establishes distinct resource landscapes that systematically select for different microbial assemblages.

### 4.2 Dietary Protein Source Impacts Microbial Diversity and Community Structure

Consistent with prior findings (Singh *et al.*, 2017; Gilbert *et al.*, 2018), plant-protein diets were associated with greater microbial alpha diversity, reflecting richer and more evenly distributed microbial communities. The Generalized Linear Mixed Model (GLMM) analysis, which modeled Shannon diversity as the dependent variable, identified protein source as the strongest predictor of alpha diversity, while protein level (% kcal), country, sample site, and host age also contributed significantly. The high conditional R² underscores the combined influence of biological and environmental variables on microbial richness (Madi *et al.*, 2023; Sweeny *et al.*, 2023). Importantly, sequencing depth did not significantly impact diversity estimates, confirming that observed patterns were biologically meaningful rather than artifacts of sampling depth (Zaheer *et al.*, 2018; Deng *et al.*, 2022).

Beta diversity analysis revealed further separation in community structure between diet groups. Principal component analysis (PCA) showed distinct clustering, supported by significant PERMANOVA and ANOSIM results. These findings align with previous studies reporting that plant-based diets promote more heterogeneous microbial communities, whereas animal-protein diets tend to support a narrower microbiome enriched in nitrogen-metabolizing taxa (Wu *et al.*, 2011; Johnson *et al.*, 2019). Together, these results suggest that protein source not only shapes microbial richness but also drives functionally distinct ecological communities.

### 4.3 Microbial Biomarkers Reflect Functional Adaptation to Protein Type

Machine learning analyses identified microbial taxa that reliably distinguished between plant- and animal-protein diets. Random Forest and LEfSe consistently highlighted *Muribaculaceae*, *Faecalibaculum*, and *Parasutterella* as biomarkers of plant-protein conditions, consistent with their known roles in saccharolytic fermentation and nitrogen salvage. *Muribaculaceae* is a dominant and functionally diverse group in murine gut ecosystems, often associated with polysaccharide utilization and propionate production (Ormerod *et al.*, 2016). Its enrichment here supports prior reports linking it to fiber-rich and lower-protein diets. *Faecalibaculum*, another SCFA-producing genus, has been implicated in carbohydrate fermentation and maintenance of gut barrier function (Zhu *et al.*, 2020). *Parasutterella*, though less frequently studied, has been linked to bile acid metabolism and appears responsive to plant-derived nitrogen sources, suggesting potential roles in amino acid salvage under lower-protein conditions.

In contrast, *Bifidobacterium*, *Streptococcus*, and *Colidextribacter* were consistently enriched in animal-protein diets, reflecting a microbial community adapted to proteolytic substrates. While *Bifidobacterium* is traditionally associated with fiber metabolism, certain species are capable of amino acid fermentation and expand in protein-rich, low-fiber contexts (Scott *et al.*, 2017). *Streptococcus* may utilize simple nitrogenous compounds and peptides from dietary or host sources, while *Colidextribacter* has recently been associated with branched-chain and aromatic amino acid fermentation in murine models.

While Random Forest provided high predictive accuracy, LEfSe identified taxa with strongest enrichment and biological effect sizes (Segata *et al.*, 2011). Together, these findings suggest that dietary protein type selectively enriches microbial lineages with distinct metabolic strategies. Our study builds on prior research by linking taxonomic signatures to predicted metabolic pathways, highlighting these taxa as potential biomarkers of diet-associated functional shifts in the gut ecosystem (Topçuoğlu *et al.*, 2020).

### 4.4 Dietary Protein Source Shapes Gut Microbial Metabolism and Functional Networks

Functional prediction using PICRUSt2 showed that plant-based and animal-based protein diets influence the metabolic potential of gut microbes in distinct ways. Diets rich in plant protein increased the abundance of pathways related to carbohydrate metabolism, nitrogen recycling, and the degradation of aromatic amino acids. These included the Entner–Doudoroff pathway, nitrate reduction, and L-tryptophan degradation, suggesting a microbial community better adapted to environments with more fiber and limited nitrogen (Fu *et al.*, 2022; Diether *et al.*, 2019). These functions are often associated with enhanced metabolic flexibility, allowing microbes to utilize a broader range of substrates, especially under conditions of nutrient limitation (Zhao *et al.*, 2019; Scott *et al.*, 2013). Additionally, pathways such as nitrate reduction provide alternative electron acceptors, while ArAA degradation contributes to detoxification and redox balancing, potentially reducing the accumulation of harmful fermentation byproducts such as ammonia, hydrogen sulfide, and p-cresol, which are more commonly associated with proteolytic metabolism (Windey *et al.*, 2012; Dai *et al.*, 2011). Thus, the enrichment of these pathways in plant-protein diets may reflect a functional shift toward a more ecologically balanced and less putrefactive microbial community. In contrast, diets enriched in animal protein promoted microbial pathways involved in proteolytic fermentation and adaptation to nitrogen-rich environments. These included creatinine degradation, propionate fermentation, and branched-chain amino acid (BCAA) biosynthesis—metabolic routes that have been linked to the generation of pro-inflammatory metabolites such as ammonia, branched-chain fatty acids, and p-cresol (Wu *et al.*, 2022; Ridlon *et al.*, 2016; Windey *et al.*, 2012). The enrichment of these functions suggests a microbial metabolic profile that favors nitrogen disposal and amino acid fermentation, processes often associated with gut barrier disruption and systemic inflammation in both animal and human studies

Network analysis further illustrated the ecological organization of these functional traits. Correlation-based mapping identified *Muribaculaceae* and *Desulfovibrio* as central nodes within plant-protein-associated subnetworks enriched in SCFA and sulfate metabolism, while *Lactobacillus* emerged as a key hub in amino acid and sulfur biosynthesis modules under animal-protein conditions (Faust & Raes, 2016; Liu *et al.*, 2021).These results suggest that dietary protein source not only alters microbial composition but reprograms functional networks, shaping key outputs like SCFA, BCFA, p-cresol, and ammonia (Neis *et al.*, 2015; Canfora *et al.*, 2019). Understanding these specific pathways is critical for linking diet to gut microbial ecology and host physiology

### 4.4 Functional Implications of Protein-Driven Microbial Shifts

Distinct microbial communities shaped by plant- and animal-based protein diets exhibited markedly different metabolic profiles, particularly in pathways involved in short-chain fatty acid (SCFA), branched-chain fatty acid (BCFA), ammonia, sulfur, and phenol metabolism. These differences reflect not only microbial adaptation to the available nutrient substrates but also carry important consequences for host metabolic health and immune regulation.

SCFA metabolism was most strongly enriched in plant-protein-fed microbiomes. Taxa such as *Muribaculaceae*, *Alistipes*, and *Bacteroides*, which were more abundant under plant-protein diets, correlated positively with glutamate degradation and lysine fermentation pathways—key routes for the production of propionate and butyrate. SCFAs are essential for maintaining intestinal barrier integrity, supporting immune regulation, and providing energy to colonocytes (Canfora *et al., 2019*; Koh *et al.*, 2016). Their anti-inflammatory and antitumor properties have been well documented, especially in the context of fiber-rich and polyphenol-containing diets (Ríos-Covián *et al.*, 2016; Louis & Flint, 2017). These findings suggest that plant-based proteins potentially promote a more resilient and protective microbial environment.

In contrast, BCFA metabolism was predominantly linked to microbiomes from animals fed animal-based protein diets. Microbes such as *Clostridium sensu stricto* 1, *Colidextribacter*, and *Lactobacillus* were associated with the fermentation of isoleucine and methionine—branched-chain and sulfur-containing amino acids that serve as substrates for proteolytic fermentation. *Clostridium sensu stricto* 1, a genus within the *Clostridia* class, harbors enzymes such as branched-chain aminotransferases and dehydrogenases that enable the conversion of BCAAs into short-chain fatty acids like isobutyrate and isovalerate, with ammonia as a byproduct via Stickland fermentation (Neis *et al.*, 2015; Bahl *et al.*, 2022). *Colidextribacter*, though recently classified, has been associated in murine studies with the enrichment of genes involved in aromatic and branched-chain amino acid degradation, suggesting its role in protein catabolism under high-protein diets (Zhao *et al.*, 2021). Likewise, certain *Lactobacillus* species can catabolize amino acids such as methionine and isoleucine under carbohydrate-limited conditions, utilizing amino acid deamination pathways to generate energy while releasing BCFA and ammonia as end-products (Pessione, 2012).

While BCFAs such as isobutyrate and isovalerate can participate in host signaling, excessive accumulation often coincides with proteolytic fermentation and mucosal stress (Smith & Macfarlane, 1997). These metabolites tend to increase in the absence of carbohydrate substrates, highlighting the trade-off between microbial fermentation strategies and host health outcomes.

Ammonia metabolism followed a similar trend. Animal-protein diets supported the enrichment of Stickland-fermenting taxa, which deaminate amino acids and increase luminal ammonia concentrations. Excessive ammonia is cytotoxic and has been associated with impaired colonocyte metabolism and barrier dysfunction (Ivnitsky *et al.*, 2022; Windey *et al.*, 2012). On the other hand, plant-protein diets appeared to favor nitrogen recycling through pathways such as glutamine synthesis. The increased abundance of *Muribaculaceae* and *Parasutterella*, both linked to ammonia assimilation, suggests a microbial adaptation that reduces nitrogen waste and recycles ammonia into biomass (Smith *et al.*, 2022; Ju *et al.*, 2019).

Sulfur metabolism further illustrates the contrast between dietary protein sources. Diets rich in animal protein increased the abundance of microbial genes involved in cysteine and methionine breakdown, which are important sources of hydrogen sulfide (H₂S). When produced in excess, H₂S can interfere with butyrate metabolism, damage cellular DNA, and weaken the intestinal barrier by disrupting tight junctions (Carbonero *et al.*, 2012; Teigen *et al.*, 2022). Microbes such as *Faecalibaculum* and *Lactobacillus*, which played central roles in sulfur-associated networks, likely contributed to elevated H₂S levels in animals consuming animal-based proteins. In contrast, the reduced presence of these pathways in mice fed plant protein may reflect a microbial community that favors alternative fermentation routes, which are less dependent on sulfur metabolism and more supportive of gut barrier function.

Finally, phenol metabolism varied depending on the source of dietary protein. *Colidextribacter*, which was more abundant in mice fed with animal-derived protein, showed strong positive correlations with tryptophan and tyrosine degradation pathways (Figure 5B). These pathways are known to produce phenolic metabolites such as *p*-cresol and indoxyl sulfate, which are classified as uremic toxins and have been associated with systemic inflammation, gut barrier dysfunction, and kidney injury (Thuy-Boun *et al.*, 2022; Wikoff *et al.*, 2009; Mishima and Abe, 2022). These compounds can disrupt gut–liver axis signaling and are increasingly recognized as biomarkers of excessive protein fermentation and microbial imbalance (Vaziri *et al.*, 2013). In contrast, *Bacteroides*, enriched under plant-based protein diets, was correlated with pathways linked to aromatic amino acid metabolism, including those involved in the production of 4-hydroxyphenylacetate (Figure 5B, 5C). This compound has been reported to exert anti-inflammatory effects and may contribute to metabolic resilience and reduced disease risk (De Filippis *et al.*, 2016; Canfora *et al.*, 2019).

Collectively, these findings emphasize that dietary protein type not only shapes microbial taxonomic profiles but also reprograms gut metabolic outputs. Plant-protein diets foster saccharolytic and nitrogen-recycling functions, while animal-protein diets enrich for proteolytic and potentially proinflammatory metabolite production. These functional shifts provide mechanistic insights into how dietary protein may influence host physiology, inflammation, and chronic disease risk.

### 4.5 Implications for Diet-Microbiome Interactions

These findings highlight the ecological and functional adaptability of the gut microbiota in response to dietary protein source. Plant-based proteins appear to support metabolic flexibility, nitrogen recycling, and anti-inflammatory capacities through enrichment of saccharolytic and amino acid degradation pathways. In contrast, animal-based proteins promote proteolytic fermentation and biosynthetic routes often associated with pro-inflammatory and potentially dysbiotic profiles. Notably, animal-protein diets are typically richer in sulfur-containing amino acids such as methionine and cysteine, which are metabolized by taxa like *Clostridium*, *Desulfovibrio*, and *Colidextribacter* into hydrogen sulfide and ammonia—metabolites associated with mucosal inflammation and compromised gut barrier integrity (Zhao *et al.*, 2021; Ridlon *et al.*, 2016).

These patterns suggest that the quality of dietary protein is potentially a key modulator of microbiome-associated metabolic health. However, caution is warranted when interpreting causality, given the potential confounding effects of co-varying dietary factors such as fiber, fat, and phytochemicals that differ across studies (Johnson *et al.*, 2019; Agus *et al.*, 2016; Holmes *et al.*, 2008). Although this meta-analysis identifies consistent associations between protein type and microbial ecology, differences in overall diet composition likely contribute to the observed microbial signatures. To isolate protein-specific effects, future studies should prioritize controlled interventions using matched isocaloric, isonitrogenous diets with standardized non-protein components.

Future work should also examine how these microbial shifts impact host physiology, particularly in relation to insulin sensitivity, lipid metabolism, immune function, and gut barrier integrity under conditions of metabolic or inflammatory stress. Integrative multi-omics approaches, including shotgun metagenomics, untargeted metabolomics, and host transcriptomics, will be essential for establishing causal links between dietary protein, microbial metabolism, and host health

### 4.6 Cross-Species Relevance and Implications for Translational Nutrition Research

Although mice and humans differ in physiology, habitual diet, and baseline microbial composition, several genera were shared across both datasets and displayed broadly similar CLR-based abundance patterns. Genera such as *Brevundimonas*, *Weissella*, *Sphingomonas*, and *Christensenella* appeared consistently across hosts, suggesting that these microbes occupy comparable ecological niches and contribute to similar metabolic functions. These observations align with prior evidence showing that specific fermentative and mucin-associated taxa maintain conserved ecological roles across mammalian gut ecosystems (Ley *et al.*, 2008; Muegge *et al.*, 2011; Nguyen *et al.*, 2015).

Several genera that responded strongly to dietary protein type in mice were also detected at measurable frequencies in human stool samples. Their presence supports the idea that metabolic pathways highlighted in murine protein-intervention models—particularly those related to amino acid fermentation and nitrogen metabolism—reflect ecological strategies not restricted to rodents. This interpretation is consistent with work demonstrating that key functional capacities of gut microbial communities, including amino acid fermentation, are frequently conserved across hosts even when taxonomic profiles differ (Turnbaugh *et al.*, 2009; Suzuki & Ley, 2020).

In this context, murine models offer a controlled and tractable framework for probing nutrient–microbe interactions with relevance for human nutrition. Their genetic uniformity, well-defined diets, and reduced environmental variability allow investigators to isolate the specific effects of dietary components in ways not feasible in free-living human populations. Such mechanistic studies have repeatedly demonstrated value for identifying microbial pathways and phenotypes that are later validated in humans (Faith *et al.*, 2014; Hildebrand *et al.*, 2013).

At the same time, the modest ΔCLR differences observed between hosts reflect expected biological variation arising from differences in anatomy, habitual diet, metabolic rate, and lifestyle. These distinctions reinforce that mice should not be viewed as direct surrogates for humans but rather as experimental systems for generating mechanistic hypotheses. The conserved genera identified here provide focused targets for future human studies incorporating strain-resolved metagenomics, metabolomics, and controlled feeding designs to evaluate how protein source shapes microbial metabolism.

### 4.7 Limitations and Recommendations for Future Research

While this meta-analysis offers valuable insights into how dietary protein sources shape the gut microbiome, several limitations should be acknowledged. First, the use of murine models limits direct translatability to humans, as differences in gut microbial composition, immune responses, and host physiology can influence microbial interactions (Nguyen *et al.*, 2015). Although our cross-species comparison highlights a subset of genera shared between mice and humans, these similarities should be interpreted cautiously, as ecological correspondence does not fully resolve species- or strain-level differences. Nevertheless, murine models remain a cornerstone in microbiome research due to their controlled genetics, defined dietary environments, and tractable experimental design. By focusing on mice, we were able to isolate the effects of dietary protein while minimizing the confounding variables, such as baseline diet variability, medication use, and lifestyle factors, that often complicate human studies. Thus, this analysis provides a foundational framework for identifying microbial and functional patterns that warrant further investigation in human cohorts.

Second, the reliance on 16S rRNA gene sequencing and functional prediction tools such as PICRUSt2 limits taxonomic resolution and cannot directly confirm gene expression or strain-level variation. While these tools are useful for comparative functional inference, they do not capture actual microbial activity. Therefore, the functional pathways identified here should be viewed as putative indicators of metabolic potential and ideally paired with direct metagenomic, metatranscriptomics, or metabolomic approaches in future work

Additionally, the absence of fecal or systemic metabolomic data precludes validation of microbial metabolic products such as short-chain fatty acids, ammonia, hydrogen sulfide, and p-cresol. As such, pathway-level inferences should be interpreted cautiously and ideally confirmed using direct metabolite profiling. Although the publicly available datasets used in this meta-analysis varied in metadata completeness, samples were retained whenever possible, and exclusions were limited only to models requiring specific metadata fields. This approach allowed for robust analysis while reducing unnecessary data loss. We compiled all available macronutrient information into **Supplementary Table 1** to document variation in diet formulation across studies. However, incomplete reporting limited our ability to incorporate these variables as covariates.

Future research should incorporate longitudinal human dietary interventions that leverage strain-resolved metagenomics, metatranscriptomics, and untargeted metabolomics to more precisely characterize microbiota function and host–microbiota interactions (Zhao *et al.*, 2019). Exploring sex-specific and age-dependent responses will also improve our understanding of how life stage and biological sex modulate microbial responses to dietary protein.

Finally, minor methodological differences, such as variation in primer pairs (e.g., 341F vs. 515F), may introduce slight biases in taxonomic profiling. However, the consistent targeting of the V3–V4 region across studies and standardized data processing pipelines likely mitigate such effects on broader community and functional trends. Together, despite these limitations, our meta-analysis offers a valuable comparative resource and highlights reproducible microbial features and functional shifts associated with dietary protein source across diverse mouse models.

## 5.0 Conclusion

This meta-analysis demonstrates that dietary protein source plays a central role in shaping gut microbial diversity, community composition, and predicted metabolic function in mice. Plant proteins enriched saccharolytic and nitrogen-recycling taxa, whereas animal proteins favored proteolytic and sulfur-metabolizing communities, aligning with distinct patterns in SCFA production, ammonia turnover, and sulfur pathways. Cross-species comparisons showed that several protein-responsive genera identified in mice also occur in humans, supporting the relevance of murine models for studying diet–microbiome interactions. These findings indicate that protein source is a modifiable dietary factor capable of altering microbial function and metabolic outputs, providing a foundation for future human interventions aimed at improving gut and metabolic health through targeted protein choices.

## Author Contributions

JHM and SA designed the study. SA, ANO and PD were involved in data screening and processing. SA and GL analyzed the data, created figures, and wrote the manuscript. ANO, JM, CKYH and JMA reviewed the manuscript. All authors participated in interpreting data, revised the manuscript, and agreed to the published version of the manuscript.

## Data Availability Statement

Data and bioinformatic workflows are publicly available here: https://github.com/deejayprof/META_ANALYSIS

## Supporting information

Meta_analysis Supplementary Tables

Meta_analysis Supplementary Figures

## Acknowledgements

We acknowledge the support provided by various R libraries that were instrumental in the analysis conducted for this study. Comprehensive details about the data, code, and resources utilized are provided in the data availability section.

## Abbreviations

PRISMA: Preferred Reporting Items for Systematic Review and Meta-Analysis
RF: Random Forest
LDA: Linear Discriminant Analysis
SCFAs: Short-Chain Fatty Acids
BCFAs: Branched-Chain Fatty Acids
ANOVA: Analysis of Variance
ROC: Receiver Operating Characteristic
AUC: Area Under the Curve
ACE: Abundance-based Coverage Estimator
PERMANOVA: Permutational Multivariate Analysis of Variance
PCoA: Principal Coordinates Analysis
GLMM: Generalized Linear Mixed Model
LEfSe: Linear Discriminant Analysis Effect Size
qPCR: Quantitative Polymerase Chain Reaction
BCAAs: Branched-Chain Amino Acids
ArAAs: Aromatic Amino Acids
ANOSIM: Analysis of Similarities
ASVs: Amplicon Sequence Variants
rRNA: Ribosomal Ribonucleic Acid
NCBI: National Center for Biotechnology Information
SRA: Sequence Read Archive
ENA: European Nucleotide Archive
QIIME2: Quantitative Insights Into Microbial Ecology 2
DADA2: Divisive Amplicon Denoising Algorithm 2
GEMs: Genome-scale metabolic models
MDA: Mean Decrease Accuracy
EAAs: Essential amino acids
PICRUST: Phylogenetic Investigation of Communities by Reconstruction of Unobserved States

## References

Rowland, I., Gibson, G., Heinken, A., Scott, K., Swann, J., Thiele, I., & Tuohy, K. (2018). Gut microbiota functions: metabolism of nutrients and other food components. European journal of nutrition, 57(1), 1–24. 10.1007/s00394-017-1445-8

Gilbert, M. S., Ijssennagger, N., Kies, A. K., & van Mil, S. W. C. (2018). Protein fermentation in the gut; implications for intestinal dysfunction in humans, pigs, and poultry. American journal of physiology. Gastrointestinal and liver physiology, 315(2), G159–G170. 10.1152/ajpgi.00319.2017

Zhao, J., Zhang, X., Liu, H., Brown, M. A., & Qiao, S. (2019). Dietary Protein and Gut Microbiota Composition and Function. Current protein & peptide science, 20(2), 145–154. 10.2174/1389203719666180514145437

Ling, ZN., Jiang, YF., Ru, JN. et al., Amino acid metabolism in health and disease. Sig Transduct Target Ther 8, 345 (2023). 10.1038/s41392-023-01569-3

Kitada, M., Ogura, Y., Monno, I., & Koya, D. (2019). The impact of dietary protein intake on longevity and metabolic health. EBioMedicine, 43, 632–640. 10.1016/j.ebiom.2019.04.005

Vinelli, V., Biscotti, P., Martini, D., Del Bo’, C., Marino, M., Meroño, T., Nikoloudaki, O., Calabrese, F. M., Turroni, S., Taverniti, V., Unión Caballero, A., Andrés-Lacueva, C., Porrini, M., Gobbetti, M., De Angelis, M., Brigidi, P., Pinart, M., Nimptsch, K., Guglielmetti, S., & Riso, P. (2022). Effects of Dietary Fibers on Short-Chain Fatty Acids and Gut Microbiota Composition in Healthy Adults: A Systematic Review. Nutrients, 14(13), 2559. 10.3390/nu14132559

den Besten, G., van Eunen, K., Groen, A. K., Venema, K., Reijngoud, D. J., & Bakker, B. M. (2013). The role of short-chain fatty acids in the interplay between diet, gut microbiota, and host energy metabolism. Journal of lipid research, 54(9), 2325–2340. 10.1194/jlr.R036012

Ríos-Covián, D., Ruas-Madiedo, P., Margolles, A., Gueimonde, M., de Los Reyes-Gavilán, C. G., & Salazar, N. (2016). Intestinal Short Chain Fatty Acids and their Link with Diet and Human Health. Frontiers in microbiology, 7, 185. 10.3389/fmicb.2016.00185

Page, M. J., McKenzie, J. E., Bossuyt, P. M., Boutron, I., Hoffmann, T. C., Mulrow, C. D., Shamseer, L., Tetzlaff, J. M., Akl, E. A., Brennan, S. E., Chou, R., Glanville, J., Grimshaw, J. M., Hróbjartsson, A., Lalu, M. M., Li, T., Loder, E. W., Mayo-Wilson, E., McDonald, S., McGuinness, L. A., … Moher, D. (2021). The PRISMA 2020 statement: an updated guideline for reporting systematic reviews. BMJ (Clinical research ed.), 372, n71. 10.1136/bmj.n71

Bolyen, E., Rideout, J. R., Dillon, M. R., Bokulich, N. A., Abnet, C. C., Al-Ghalith, G. A., Alexander, H., Alm, E. J., Arumugam, M., Asnicar, F., Bai, Y., Bisanz, J. E., Bittinger, K., Brejnrod, A., Brislawn, C. J., Brown, C. T., Callahan, B. J., Caraballo-Rodríguez, A. M., Chase, J., Cope, E. K., … Caporaso, J. G. (2019). Reproducible, interactive, scalable and extensible microbiome data science using QIIME 2. Nature biotechnology, 37(8), 852–857. 10.1038/s41587-019-0209-9

Ziemski, M., Adamov, A., Kim, L., Flörl, L., & Bokulich, N. A. (2022). Reproducible acquisition, management and meta-analysis of nucleotide sequence (meta)data using q2-fondue. *Bioinformatics (Oxford*, England), 38(22), 5081–5091. 10.1093/bioinformatics/btac639

Callahan, B. J., McMurdie, P. J., Rosen, M. J., Han, A. W., Johnson, A. J., & Holmes, S. P. (2016). DADA2: High-resolution sample inference from Illumina amplicon data. Nature methods, 13(7), 581–583. 10.1038/nmeth.3869

Bokulich, N. A., Kaehler, B. D., Rideout, J. R., Dillon, M., Bolyen, E., Knight, R., Huttley, G. A., & Gregory Caporaso, J. (2018). Optimizing taxonomic classification of marker-gene amplicon sequences with QIIME 2’s q2-feature-classifier plugin. Microbiome, 6(1), 90. 10.1186/s40168-018-0470-z

Wen, T., Niu, G., Chen, T., Shen, Q., Yuan, J., & Liu, Y. X. (2023). The best practice for microbiome analysis using R. Protein & cell, 14(10), 713–725. 10.1093/procel/pwad024

McMurdie, P. J., & Holmes, S. (2012). Phyloseq: a bioconductor package for handling and analysis of high-throughput phylogenetic sequence data. *Pacific Symposium on Biocomputing*. Pacific Symposium on Biocomputing, 235–246.

Samuthpongtorn, C., Nopsopon, T., & Pongpirul, K. (2022). Gut microbiome diversity measures for metabolic conditions: a systematic scoping review. Journal of Physiological and Biomedical Sciences, 33(2). retrieved from https://li01.tci-thaijo.org/index.php/j-pbs/article/view/254538

Manandhar, I., Alimadadi, A., Aryal, S., Munroe, P. B., Joe, B., & Cheng, X. (2021). Gut microbiome-based supervised machine learning for clinical diagnosis of inflammatory bowel diseases. American journal of physiology. Gastrointestinal and liver physiology, 320(3), G328–G337. 10.1152/ajpgi.00360.2020

Suzuki, T. A., & Ley, R. E. (2020). The role of the microbiota in human genetic adaptation. *Science (New York*, N.Y.), 370(6521), eaaz6827. 10.1126/science.aaz6827

Walters, W. A., Xu, Z., & Knight, R. (2014). Meta-analyses of human gut microbes associated with obesity and IBD. FEBS letters, 588(22), 4223–4233. 10.1016/j.febslet.2014.09.039

De Filippis, F., Pellegrini, N., Vannini, L., Jeffery, I. B., La Storia, A., Laghi, L., Serrazanetti, D. I., Di Cagno, R., Ferrocino, I., Lazzi, C., Turroni, S., Cocolin, L., Brigidi, P., Neviani, E., Gobbetti, M., O’Toole, P. W., & Ercolini, D. (2016). High-level adherence to a Mediterranean diet beneficially impacts the gut microbiota and associated metabolome. Gut, 65(11), 1812–1821. 10.1136/gutjnl-2015-309957

Canfora, E. E., Meex, R. C. R., Venema, K., & Blaak, E. E. (2019). Gut microbial metabolites in obesity, NAFLD and T2DM. *Nature reviews. Endocrinology*, *15*(5), 261–273. 10.1038/s41574-019-0156-z

Sweeny, A. R., Lemon, H., Ibrahim, A., Watt, K. A., Wilson, K., Childs, D. Z., Nussey, D. H., Free, A., & McNally, L. (2023). A mixed-model approach for estimating drivers of microbiota community composition and differential taxonomic abundance. mSystems, 8(4), e0004023. 10.1128/msystems.00040-23

Madi, N., Chen, D., Wolff, R., Shapiro, B. J., & Garud, N. R. (2023). Community diversity is associated with intra-species genetic diversity and gene loss in the human gut microbiome. eLife, 12, e78530. 10.7554/eLife.78530

Zaheer, R., Noyes, N., Ortega Polo, R., Cook, S. R., Marinier, E., Van Domselaar, G., Belk, K. E., Morley, P. S., & McAllister, T. A. (2018). Impact of sequencing depth on the characterization of the microbiome and resistome. Scientific reports, 8(1), 5890. 10.1038/s41598-018-24280-8

Deng, X. L., Frandsen, P. B., Dikow, R. B., Favre, A., Shah, D. N., Shah, R. D. T., Schneider, J. V., Heckenhauer, J., & Pauls, S. U. (2022). The impact of sequencing depth and relatedness of the reference genome in population genomic studies: A case study with two caddisfly species (Trichoptera, Rhyacophilidae, Himalopsyche). Ecology and evolution, 12(12), e9583. 10.1002/ece3.9583

Li, Z., Zhou, J., Liang, H., Ye, L., Lan, L., Lu, F., Wang, Q., Lei, T., Yang, X., Cui, P., & Huang, J. (2022). Differences in Alpha Diversity of Gut Microbiota in Neurological Diseases. Frontiers in neuroscience, 16, 879318. 10.3389/fnins.2022.879318

Singh, R. K., Chang, H. W., Yan, D., Lee, K. M., Ucmak, D., Wong, K., Abrouk, M., Farahnik, B., Nakamura, M., Zhu, T. H., Bhutani, T., & Liao, W. (2017). Influence of diet on the gut microbiome and implications for human health. Journal of translational medicine, 15(1), 73. 10.1186/s12967-017-1175-y

Gilbert, J. A., Blaser, M. J., Caporaso, J. G., Jansson, J. K., Lynch, S. V., & Knight, R. (2018). Current understanding of the human microbiome. Nature medicine, 24(4), 392–400. 10.1038/nm.4517

Segata, N., Izard, J., Waldron, L., Gevers, D., Miropolsky, L., Garrett, W. S., & Huttenhower, C. (2011). Metagenomic biomarker discovery and explanation. Genome biology, 12(6), R60. 10.1186/gb-2011-12-6-r60

Ju, T., Kong, J. Y., Stothard, P., & Willing, B. P. (2019). Defining the role of *Parasutterella*, a previously uncharacterized member of the core gut microbiota. The ISME journal, 13(6), 1520–1534. 10.1038/s41396-019-0364-5

Johnson, A. J., Vangay, P., Al-Ghalith, G. A., et al., (2019). Daily sampling reveals personalized diet-microbiome associations in humans. Cell Host & Microbe, 25(6), 789–802.e5. 10.1016/j.chom.2019.05.005

Tang, Z. Z., Chen, G., & Alekseyenko, A. V. (2016). PERMANOVA-S: association test for microbial community composition that accommodates confounders and multiple distances. Bioinformatics (Oxford, England), 32(17), 2618–2625. 10.1093/bioinformatics/btw311

Scott, K. P., Gratz, S. W., Sheridan, P. O., Flint, H. J., & Duncan, S. H. (2013). The influence of diet on the gut microbiota. Pharmacological research, 69(1), 52–60. 10.1016/j.phrs.2012.10.020

Wu, G. D., Chen, J., Hoffmann, C., Bittinger, K., Chen, Y. Y., Keilbaugh, S. A., Bewtra, M., Knights, D., Walters, W. A., Knight, R., Sinha, R., Gilroy, E., Gupta, K., Baldassano, R., Nessel, L., Li, H., Bushman, F. D., & Lewis, J. D. (2011). Linking long-term dietary patterns with gut microbial enterotypes. *Science (New York*, N.Y.), 334(6052), 105–108. 10.1126/science.1208344

Beaumont, M., Portune, K. J., Steuer, N., Lan, A., Cerrudo, V., Audebert, M., Dumont, F., Mancano, G., Khodorova, N., Andriamihaja, M., Airinei, G., Tomé, D., Benamouzig, R., Davila, A. M., Claus, S. P., Sanz, Y., & Blachier, F. (2017). Quantity and source of dietary protein influence metabolite production by gut microbiota and rectal mucosa gene expression: a randomized, parallel, double-blind trial in overweight humans. The American journal of clinical nutrition, 106(4), 1005–1019. 10.3945/ajcn.117.158816

Rooks, M. G., & Garrett, W. S. (2016). Gut microbiota, metabolites and host immunity. Nature reviews. Immunology, 16(6), 341–352. 10.1038/nri.2016.42

Topçuoğlu BD, Lesniak NA, Ruffin MT, Wiens J, Schloss PD. 2020. A Framework for Effective Application of Machine Learning to Microbiome-Based Classification Problems. mBio11:10.1128/mbio.00434-20. 10.1128/mbio.00434-20

Ridlon, J. M., Harris, S. C., Bhowmik, S., Kang, D. J., & Hylemon, P. B. (2016). Consequences of bile salt biotransformations by intestinal bacteria. Gut microbes, 7(1), 22–39. 10.1080/19490976.2015.1127483

Tan, J., Ni, D., Taitz, J., Pinget, G. V., Read, M., Senior, A., Wali, J. A., Elnour, R., Shanahan, E., Wu, H., Chadban, S. J., Nanan, R., King, N. J. C., Grau, G. E., Simpson, S. J., & Macia, L. (2022). Dietary protein increases T-cell-independent sIgA production through changes in gut microbiota-derived extracellular vesicles. Nature communications, 13(1), 4336. 10.1038/s41467-022-31761-y

Masuoka, H., Suda, W., Tomitsuka, E., Shindo, C., Takayasu, L., Horwood, P., Greenhill, A. R., Hattori, M., Umezaki, M., & Hirayama, K. (2020). The influences of low protein diet on the intestinal microbiota of mice. Scientific reports, 10(1), 17077. 10.1038/s41598-020-74122-9

Yin, Y., Cai, J., Zhou, L., Xing, L., & Zhang, W. (2022). Dietary oxidized beef protein alters gut microbiota and induces colonic inflammatory damage in C57BL/6 mice. Frontiers in nutrition, 9, 980204. 10.3389/fnut.2022.980204

Zhu, Y., He, H., Tang, Y., Peng, Y., Hu, P., Sun, W., Liu, P., Jin, M., & Xu, X. (2022). Reno-Protective Effect of Low Protein Diet Supplemented With α-Ketoacid Through Gut Microbiota and Fecal Metabolism in 5/6 Nephrectomized Mice. Frontiers in nutrition, 9, 889131. 10.3389/fnut.2022.889131

Xie, Y., Wang, C., Zhao, D., Zhou, G., & Li, C. (2020). Processing Method Altered Mouse Intestinal Morphology and Microbial Composition by Affecting Digestion of Meat Proteins. Frontiers in microbiology, 11, 511. 10.3389/fmicb.2020.00511

Choi, B. S., Daniel, N., Houde, V. P., Ouellette, A., Marcotte, B., Varin, T. V., Vors, C., Feutry, P., Ilkayeva, O., Ståhlman, M., St-Pierre, P., Bäckhed, F., Tremblay, A., White, P. J., & Marette, A. (2021). Feeding diversified protein sources exacerbates hepatic insulin resistance via increased gut microbial branched-chain fatty acids and mTORC1 signaling in obese mice. Nature communications, 12(1), 3377. 10.1038/s41467-021-23782-w

Ijaz, M. U., Ahmad, M. I., Hussain, M., Khan, I. A., Zhao, D., & Li, C. (2020). Meat Protein in High-Fat Diet Induces Adipogensis and Dyslipidemia by Altering Gut Microbiota and Endocannabinoid Dysregulation in the Adipose Tissue of Mice. Journal of agricultural and food chemistry, 68(13), 3933–3946. 10.1021/acs.jafc.0c00017

Zhong, W., Wang, H., Yang, Y., Zhang, Y., Lai, H., Cheng, Y., Yu, H., Feng, N., Huang, R., Liu, S., Yang, S., Hao, T., Zhang, B., Ying, H., Zhang, F., Guo, F., & Zhai, Q. (2022). High-protein diet prevents fat mass increase after dieting by counteracting Lactobacillus-enhanced lipid absorption. Nature metabolism, 4(12), 1713–1731. 10.1038/s42255-022-00687-6

Zhang, R., Mu, H., Li, Z., Zeng, J., Zhou, Q., Li, H., Wang, S., Li, X., Zhao, X., Sun, L., Chen, W., Dong, J., & Yang, R. (2022). Oral administration of branched-chain amino acids ameliorates high-fat diet-induced metabolic-associated fatty liver disease *via* gut microbiota-associated mechanisms. Frontiers in microbiology, 13, 920277. 10.3389/fmicb.2022.920277

Ruocco, C., Ragni, M., Rossi, F., Carullo, P., Ghini, V., Piscitelli, F., Cutignano, A., Manzo, E., Ioris, R. M., Bontems, F., Tedesco, L., Greco, C. M., Pino, A., Severi, I., Liu, D., Ceddia, R. P., Ponzoni, L., Tenori, L., Rizzetto, L., Scholz, M., … Nisoli, E. (2020). Manipulation of Dietary Amino Acids Prevents and Reverses Obesity in Mice Through Multiple Mechanisms That Modulate Energy Homeostasis. Diabetes, 69(11), 2324–2339. 10.2337/db20-0489

Douglas, G. M., Maffei, V. J., Zaneveld, J., Yurgel, S. N., Brown, J. R., Taylor, C. M., Huttenhower, C., & Langille, M. G. I. (2020). PICRUSt2 for prediction of metagenome functions. Nature Biotechnology, 38(6), 685–688. 10.1038/s41587-020-0548-6

Mallick, H., Rahnavard, A., McIver, L. J., Ma, S., Zhang, Y., Nguyen, L. H., Tickle, T. L., Weingart, G., Ren, B., Schwager, E. H., Thompson, K. N., Wilkinson, J. E., Subramanian, A., Lu, Y., Waldron, L., Paulson, J. N., & Huttenhower, C. (2021). Multivariable association discovery in population-scale meta-omics studies. PLOS Computational Biology, 17(11), e1009442. 10.1371/journal.pcbi.1009442

Faust, K., & Raes, J. (2016). CoNet app: Inference of biological association networks using Cytoscape. F1000Research, 5, 1519. 10.12688/f1000research.9050.2

Kuntal, B. K., Mande, S. S., & Dutta, A. (2019). CompNet: A GUI based tool for comparison of multiple biological interaction networks. BMC Bioinformatics, 20(1), 185. 10.1186/s12859-019-2721-3

Liu, Y., Amit, G., Zhao, X., Wu, N., Li, D., & Bashan, A. (2023). Individualized network analysis reveals a link between the gut microbiome, diet intervention and Gestational Diabetes Mellitus. PLoS computational biology, 19(6), e1011193. 10.1371/journal.pcbi.1011193

Liu, F., Li, Z., Wang, X., Xue, C., Tang, Q., & Li, R. W. (2019). Microbial Co-Occurrence Patterns and Keystone Species in the Gut Microbial Community of Mice in Response to Stress and Chondroitin Sulfate Disaccharide. International journal of molecular sciences, 20(9), 2130. 10.3390/ijms20092130

Windey, K., De Preter, V., & Verbeke, K. (2012). Relevance of protein fermentation to gut health. Molecular nutrition & food research, 56(1), 184–196. 10.1002/mnfr.201100542

Rutherfurd, S. M., Fanning, A. C., Miller, B. J., & Moughan, P. J. (2015). Protein digestibility-corrected amino acid scores and digestible indispensable amino acid scores differentially describe protein quality in growing male rats. The Journal of nutrition, 145(2), 372–379. 10.3945/jn.114.195438

Phillips, S. M., Chevalier, S., & Leidy, H. J. (2016). Protein “requirements” beyond the RDA: implications for optimizing health. Applied physiology, nutrition, and metabolism = Physiologie appliquee, nutrition et metabolisme, 41(5), 565–572. 10.1139/apnm-2015-0550

Zhou, W., Sailani, M. R., Contrepois, K., Zhou, Y., Ahadi, S., Leopold, S. R., Zhang, M. J., Rao, V., Avina, M., Mishra, T., Johnson, J., Lee-McMullen, B., Chen, S., Metwally, A. A., Tran, T. D. B., Nguyen, H., Zhou, X., Albright, B., Hong, B. Y., Petersen, L., … Snyder, M. (2019). Longitudinal multi-omics of host-microbe dynamics in prediabetes. Nature, 569(7758), 663–671. 10.1038/s41586-019-1236-x

Yassour, M., Vatanen, T., Siljander, H., Hämäläinen, A. M., Härkönen, T., Ryhänen, S. J., Franzosa, E. A., Vlamakis, H., Huttenhower, C., Gevers, D., Lander, E. S., Knip, M., DIABIMMUNE Study Group, & Xavier, R. J. (2016). Natural history of the infant gut microbiome and impact of antibiotic treatment on bacterial strain diversity and stability. Science translational medicine, 8(343), 343ra81. 10.1126/scitranslmed.aad0917

Ridlon, J. M., Wolf, P. G., & Gaskins, H. R. (2016). Taurocholic acid metabolism by gut microbes and colon cancer. Gut microbes, 7(3), 201–215. 10.1080/19490976.2016.1150414

Wu, S., Bhat, Z. F., Gounder, R. S., Mohamed Ahmed, I. A., Al-Juhaimi, F. Y., Ding, Y., & Bekhit, A. E. A. (2022). Effect of Dietary Protein and Processing on Gut Microbiota-A Systematic Review. Nutrients, 14(3), 453. 10.3390/nu14030453

Blakeley-Ruiz, J. A., Bartlett, A., McMillan, A. S., Awan, A., Walsh, M. V., Meyerhoffer, A. K., Vintila, S., Maier, J. L., Richie, T. G., Theriot, C. M., & Kleiner, M. (2025). Dietary protein source alters gut microbiota composition and function. The ISME journal, 19(1), wraf048. 10.1093/ismejo/wraf048

Fu, J., Zheng, Y., Gao, Y., & Xu, W. (2022). Dietary Fiber Intake and Gut Microbiota in Human Health. Microorganisms, 10(12), 2507. 10.3390/microorganisms10122507

Diether, N. E., & Willing, B. P. (2019). Microbial Fermentation of Dietary Protein: An Important Factor in Diet⁻Microbe⁻Host Interaction. Microorganisms, 7(1), 19. 10.3390/microorganisms7010019

Ivnitsky, J. J., Schäfer, T. V., Rejniuk, V. L., & Vakunenkova, O. A. (2022). Secondary Dysfunction of the Intestinal Barrier in the Pathogenesis of Complications of Acute Poisoning. Journal of evolutionary biochemistry and physiology, 58(4), 1075–1098. 10.1134/S0022093022040123

Carbonero, F., Benefiel, A. C., Alizadeh-Ghamsari, A. H., & Gaskins, H. R. (2012). Microbial pathways in colonic sulfur metabolism and links with health and disease. Frontiers in physiology, 3, 448. 10.3389/fphys.2012.00448

Koh, A., De Vadder, F., Kovatcheva-Datchary, P., & Bäckhed, F. (2016). From dietary fiber to host physiology: Short-chain fatty acids as key bacterial metabolites. Cell, 165(6), 1332–1345. 10.1016/j.cell.2016.05.041

Ríos-Covián, D., Ruas-Madiedo, P., Margolles, A., Gueimonde, M., de los Reyes-Gavilán, C. G., & Salazar, N. (2016). Intestinal short chain fatty acids and their link with diet and human health. Frontiers in Microbiology, 7, 185. 10.3389/fmicb.2016.00185

Louis, P., & Flint, H. J. (2017). Formation of propionate and butyrate by the human colonic microbiota. Environmental Microbiology, 19(1), 29–41. 10.1111/1462-2920.13589

Zhao, L., Zhang, F., Ding, X., Wu, G., & Wu, C. (2021). Gut bacteria selectively promoted by dietary fibers alleviate type 2 diabetes. Science, 359(6380), 1151–1156. 10.1126/science.aao5771

Neis, E. P. J. G., Dejong, C. H. C., & Rensen, S. S. (2015). The role of microbial amino acid metabolism in host metabolism. Nutrients, 7(4), 2930–2946. 10.3390/nu7042930

Smith, E. A., & Macfarlane, G. T. (1997). Formation of phenolic and indolic compounds by anaerobic bacteria in the human large intestine. Microbial Ecology, 33(3), 180–188. 10.1007/s002489900048

Windey, K., De Preter, V., & Verbeke, K. (2012). Relevance of protein fermentation to gut health. Molecular Nutrition & Food Research, 56(1), 184–196. 10.1002/mnfr.201100507

Teigen, L., Biruete, A., & Khoruts, A. (2023). Impact of diet on hydrogen sulfide production: implications for gut health. Current opinion in clinical nutrition and metabolic care, 26(1), 55–58. 10.1097/MCO.0000000000000881

Thuy-Boun, P. S., Wang, A. Y., Crissien-Martinez, A., Xu, J. H., Chatterjee, S., Stupp, G. S., Su, A. I., Coyle, W. J., & Wolan, D. W. (2022). Quantitative Metaproteomics and Activity-based Protein Profiling of Patient Fecal Microbiome Identifies Host and Microbial Serine-type Endopeptidase Activity Associated With Ulcerative Colitis. Molecular & cellular proteomics : MCP, 21(3), 100197. 10.1016/j.mcpro.2022.100197

Mishima, E., & Abe, T. (2022). Gut Microbiota Dynamics and Uremic Toxins. Toxins, 14(2), 146. 10.3390/toxins14020146

Vaziri, N. D., Wong, J., Pahl, M., Piceno, Y. M., Yuan, J., DeSantis, T. Z., Ni, Z., Nguyen, T. H., & Andersen, G. L. (2013). Chronic kidney disease alters intestinal microbial flora. Kidney international, 83(2), 308–315. 10.1038/ki.2012.345

Parada, A. E., Needham, D. M., & Fuhrman, J. A. (2016). Every base matters: assessing small subunit rRNA primers for marine microbiomes with mock communities, time series and global field samples. Environmental microbiology, 18(5), 1403–1414. 10.1111/1462-2920.13023

Klindworth, A., Pruesse, E., Schweer, T., Peplies, J., Quast, C., Horn, M., & Glöckner, F. O. (2013). Evaluation of general 16S ribosomal RNA gene PCR primers for classical and next-generation sequencing-based diversity studies. Nucleic acids research, 41(1), e1. 10.1093/nar/gks808

Guoyao Wu. Amino acids in nutrition, health, and disease. Front. Biosci. (Landmark Ed) 2021, 26(12), 1386–1392. 10.52586/5032

Holmes, E., Loo, R. L., Stamler, J., Bictash, M., Yap, I. K., Chan, Q., Ebbels, T., De Iorio, M., Brown, I. J., Veselkov, K. A., Daviglus, M. L., Kesteloot, H., Ueshima, H., Zhao, L., Nicholson, J. K., & Elliott, P. (2008). Human metabolic phenotype diversity and its association with diet and blood pressure. Nature, 453(7193), 396–400. 10.1038/nature06882

Agus, A., Denizot, J., Thévenot, J., Martinez-Medina, M., Massier, S., Sauvanet, P., Bernalier-Donadille, A., Denis, S., Hofman, P., Bonnet, R., Billard, E., & Barnich, N. (2016). Western diet induces a shift in microbiota composition enhancing susceptibility to Adherent-Invasive E. coli infection and intestinal inflammation. Scientific reports, 6, 19032. 10.1038/srep19032

Johnson, A. J., Vangay, P., Al-Ghalith, G. A., Hillmann, B. M., Ward, T. L., Shields-Cutler, R. R., Kim, A. D., Shmagel, A. K., Syed, A. N., Personalized Microbiome Class Students, Walter, J., Menon, R., Koecher, K., & Knights, D. (2019). Daily Sampling Reveals Personalized Diet-Microbiome Associations in Humans. Cell host & microbe, 25(6), 789–802.e5. 10.1016/j.chom.2019.05.005

Gloor, G. B., Macklaim, J. M., Pawlowsky-Glahn, V., & Egozcue, J. J. (2017). Microbiome Datasets Are Compositional: And This Is Not Optional. Frontiers in microbiology, 8, 2224. 10.3389/fmicb.2017.02224

Hildebrand, F., Nguyen, T. L., Brinkman, B., Yunta, R. G., Cauwe, B., Vandenabeele, P., Liston, A., & Raes, J. (2013). Inflammation-associated enterotypes, host genotype, cage and inter-individual effects drive gut microbiota variation in common laboratory mice. Genome biology, 14(1), R4. 10.1186/gb-2013-14-1-r4

Nguyen, T. L., Vieira-Silva, S., Liston, A., & Raes, J. (2015). How informative is the mouse for human gut microbiota research? Disease models & mechanisms, 8(1), 1–16. 10.1242/dmm.017400

Adejumo SA, Oli AN, Rowaiye AB, Igbokwe NH, Ezejiegu CK, Yahaya ZS. Immunomodulatory Benefits of Probiotic Bacteria: A Review of Evidence. OBM Genetics 2023; 7(4): 206; doi:10.21926/obm.genet.2304206.

Turnbaugh, P. J., Ridaura, V. K., Faith, J. J., Rey, F. E., Knight, R., & Gordon, J. I. (2009). The effect of diet on the human gut microbiome: a metagenomic analysis in humanized gnotobiotic mice. Science translational medicine, 1(6), 6ra14. 10.1126/scitranslmed.3000322

Quinn, T. P., Erb, I., Richardson, M. F., & Crowley, T. M. (2018). Understanding sequencing data as compositions: an outlook and review. *Bioinformatics (Oxford*, England), 34(16), 2870–2878. 10.1093/bioinformatics/bty175

McDonald, D., Hyde, E., Debelius, J. W., Morton, J. T., Gonzalez, A., Ackermann, G., Aksenov, A. A., Behsaz, B., Brennan, C., Chen, Y., DeRight Goldasich, L., Dorrestein, P. C., Dunn, R. R., Fahimipour, A. K., Gaffney, J., Gilbert, J. A., Gogul, G., Green, J. L., Hugenholtz, P., Humphrey, G., et al. (2018). American Gut: an Open Platform for Citizen Science Microbiome Research. mSystems, 3(3), e00031–18. 10.1128/mSystems.00031-18

Lahti, L., & Shetty, S. (2017). microbiome R package. Bioconductor. 10.18129/B9.bioc.microbiome

Ley, R. E., Hamady, M., Lozupone, C., Turnbaugh, P. J., Ramey, R. R., Bircher, J. S., Schlegel, M. L., Tucker, T. A., Schrenzel, M. D., Knight, R., & Gordon, J. I. (2008). Evolution of mammals and their gut microbes. *Science (New York*, N.Y.), 320(5883), 1647–1651. 10.1126/science.1155725

Faith JJ, Ahern PP, Ridaura VK, Cheng J, Gordon JI. Identifying gut microbe-host phenotype relationships using combinatorial communities in gnotobiotic mice. Sci Transl Med. 2014 Jan 22;6(220):220ra11. doi: 10.1126/scitranslmed.3008051. PMID: 24452263; PMCID: PMC3973144.

Muegge, B. D., Kuczynski, J., Knights, D., Clemente, J. C., González, A., Fontana, L., Henrissat, B., Knight, R., & Gordon, J. I. (2011). Diet drives convergence in gut microbiome functions across mammalian phylogeny and within humans. *Science (New York*, N.Y.), 332(6032), 970–974. 10.1126/science.1198719

