## Supplementary material for "Dietary Protein Source Shapes Gut Microbial Structure and Predicted Function: A Meta-Analysis with Machine Learning": Meta_analysis Supplementary Tables

**Supplementary Table 1. 16S rRNA Gene Primers and Regions used in included studies**

| Country, Year, First Author | Bioproject accession | Target Region | Forward Primer | Reverse Primer |
| --- | --- | --- | --- | --- |
| Australia, 2022  Jian Tan | PRJEB39583 | V3-V4 | 515F (5′-GTGCCAGCMGCCGCGG-3′) | 806R (GGACTACHVGGGTWTCTAAT) |
| Japan, 2020  Hiroaki Masuoka | PRJDB8898 | V1- V2 | 27F (5′-AGAGTTTGATCMTGGCTCAG-3′) | 338R (TGCTGCCTCCCGTAGGAGT) |
| China, 2022  Yantao Yin | PRJNA872483 | V3-V4 | 515F (5′-GTGCCAGCMGCCGCGG-3′) | 806R (GGACTACHVGGGTWTCTAAT) |
| China, 2022  Yifan Zhu | PRJNA810785 | V3-V4 | 338F (5′-ACTCCTACGGGAGGCAGCAG-3′) | 806R (GGACTACHVGGGTWTCTAAT) |
| China, 2020  Yunting Xie | PRJNA607338 | V3-V4 | 515F (5′-GTGCCAGCMGCCGCGG-3′) | 806R (GGACTACHVGGGTWTCTAAT) |
| Canada, 2021  Béatrice S.-Y. Choi | PRJEB37442 | V3-V4 | 515F (5′-GTGCCAGCMGCCGCGG-3′) | 806R (GGACTACHVGGGTWTCTAAT) |
| China, 2020  Muhammad Umair Ijaz | PRJNA548036 | V3-V4 | 515F (5′-GTGCCAGCMGCCGCGG-3′) | 806R (GGACTACHVGGGTWTCTAAT) |
| China, 2022  Wuling Zhong | PRJNA868153 | V3-V4 | 515F (5′-GTGCCAGCMGCCGCGG-3′) | 806R (GGACTACHVGGGTWTCTAAT) |
| China, 2022  Ranran Zhang | PRJNA852193 | V3-V4 | 338F (ACTCCTACGGGAGGCAGCAG) | 806R (GGACTACHVGGGTWTCTAAT) |
| Italy, 2020  Chiara Ruocco | PRJEB25686 | V3-V4 | 515F (5′-GTGCCAGCMGCCGCGG-3′) | 806R (GGACTACHVGGGTWTCTAAT) |

**Supplementary Table 2. Macronutrient composition (energy-based %kcal) for all rodent diets included in the meta-analysis.**

| Bioproject accession | Protein (%kcal) | Fat (%kcal) | Carbohydrate (incl. fiber) (%kcal) | Protein type | Country, Year, First Author |
| --- | --- | --- | --- | --- | --- |
| PRJEB39583 | 60 | 20 | 20 | Casein | Australia, 2022  Jian Tan |
| PRJDB8898 | 3 | 7 | 81.6 | Casein | Japan, 2020  Hiroaki Masuoka |
| PRJDB8898 | 12 | 7 | 71.5 | Casein | Japan, 2020  Hiroaki Masuoka |
| PRJNA872483 | 17.86 | 2.9 | 62.95 | Oxidized beef | China, 2022  Yantao Yin |
| PRJNA810785 | 5 | 4-7 | 70-80 | Wheat starch–based low-protein diet | China, 2022  Yifan Zhu |
| PRJNA607338 | 20 | 7.0 | 62.95 | Casein, Pork, Sausage, Soy protein | China, 2020  Yunting Xie |
| PRJEB37442 | 15 | 10 | 75 | Casein | Canada, 2021  Béatrice S.-Y. Choi |
| PRJNA548036 | 20 | 10.1 | 69.9 | Beef, Casein, Chicken, Pork | China, 2020  Muhammad Umair Ijaz |
| PRJNA868153 | 61.5 | 15.9 | 22.5 | Casein | China, 2022  Wuling Zhong |
| PRJNA852193 | 25 | 12 | 63 | Branched-chain amino acids | China, 2022  Ranran Zhang |
| PRJEB25686 | 20 | 10 | 70 | Essential amino acids, Casein | Italy, 2020  Chiara Ruocco |

**Supplementary Table 3: Core microbiota based on protein source at the Phylum level**

| Phyla | Animal Protein (%) | Phyla | Plant Protein (%) |
| --- | --- | --- | --- |
| *Firmicutes* | 63.34 | *Firmicutes* | 47.25 |
| *Bacteroidota* | 19.96 | *Bacteroidota* | 23.45 |
| *Proteobacteria* | 4.78 | *Proteobacteria* | 10.21 |
| *Actinobacteriota* | 4.16 | *Actinobacteriota* | 3.82 |
| *Verrucomicrobiota* | 1.43 | *Acidobacteriota* | 3.19 |
| *Desulfobacterota* | 1.03 | *Chloroflexi* | 1.61 |
| *Cyanobacteria* | 0.68 | *Cyanobacteria* | 1.30 |
| *Deferribacterota* | 0.60 | *Verrucomicrobiota* | 0.97 |
| *Patescibacteria* | 0.55 | *Desulfobacterota* | 0.85 |
| *Campylobacterota* | 0.39 | *Campylobacterota* | 0.66 |
| *Others* | 3.07 | Others | 6.68 |

**Supplementary Table 4: General Linear Mixed-Effects model comparison**

| Metric | Model 1 | Model 4 | Model 5 | Model 6 | Model 2 | Model 7 | Model 3 |
| --- | --- | --- | --- | --- | --- | --- | --- |
| Number of Parameters (npar) | 4 | 5 | 6 | 7 | 8 | 8 | 9 |
| Akaike Information Criterion (AIC) | 456.49 | 417.20 | 414.68 | 250.95 | 356.27 | 252.88 | 275.84 |
| Bayesian Information Criterion (BIC) | 469.15 | 433.02 | 433.67 | 273.11 | 381.59 | 278.20 | 304.32 |
| Log-Likelihood (logLik) | -224.25 | -203.60 | -201.34 | -118.48 | -170.13 | -118.44 | -128.92 |
| Deviance | 448.49 | 407.20 | 402.68 | 236.95 | 340.27 | 236.88 | 257.84 |
| Chi-squared Test (Chisq) | - | 41.29 | 4.52 | 165.73 | - | 103.39 | - |
| Degrees of Freedom (Df) | - | 1 | 1 | 1 | 1 | 0 | 1 |
| p-value (Pr(>Chisq)) | - | 1.31e-10 | 0.03356 | < 2.2e-16 | 1 | - | 1 |

**Models:**

M1: Shannon ~ Protein_source + (1 | Isolation_source)

M4: Shannon ~ Protein_source + Protein_level + (1 | Isolation_source)

M5: Shannon ~ Protein_source + Protein_level + Age_in_weeks + (1 | Isolation_source)

M6: Shannon ~ Protein_source + Protein_level + (1 | Country) + (1 | Age_in_weeks) + (1 | Isolation_source)

M2: Shannon ~ Protein_source + Protein_level + Country + (1 | Isolation_source)

M7: Shannon ~ Protein_source * Protein_level + (1 | Country) + (1 | Age_in_weeks) + (1 | Isolation_source)

M3: Shannon ~ Protein_source + Protein_level + Age_in_weeks + Country + (1 | Isolation_source)

**Fixed Effects Estimates:**

Intercept: 0.33899, Protein_level: 0.40397, Protein_source: 1.25097, Age_in_weeks: 0.30001, Country: -0.55194

**Supplementary Table 5: Random Forest Mean Decrease Accuracy Analysis of Top 10 Taxa**

| Mean Decrease Accuracy | Kingdom | Phylum | Class | Order | Family | Genus |
| --- | --- | --- | --- | --- | --- | --- |
| 5.344502894 | Bacteria | *Bacteroidota* | *Bacilli* | *Lactobacillales* | *Streptococcaceae* | *Lactococcus* |
| 5.094711866 | Bacteria | *Firmicutes* | *Bacilli* | *Lactobacillales* | *Streptococcaceae* | *Streptococcus* |
| 4.707552935 | Bacteria | *Firmicutes* | *Clostridia* | *Peptococcales* | *Peptococcaceae* | *Peptococcus* |
| 4.432882205 | Bacteria | *Firmicutes* | *Bacilli* | *Erysipelotrichales* | *Erysipelotrichaceae* | *Faecalibaculum* |
| 4.437837578 | Bacteria | *Actinobacteriota* | *Coriobacteriia* | *Coriobacteriales* | *Atopobiaceae* | *Coriobacteriaceae_UCG-002* |
| 4.349395422 | Bacteria | *Firmicutes* | *Bacilli* | *Lactobacillales* | *Streptococcaceae* | *Streptococcus* |
| 4.558971351 | Bacteria | *Actinobacteriota* | *Actinobacteria* | *Bifidobacteriales* | *Bifidobacteriaceae* | *Bifidobacterium* |
| 4.465968838 | Bacteria | *Proteobacteria* | *Gammaproteobacteria* | *Burkholderiales* | *Sutterellaceae* | *Parasutterella* |
| 4.10452852 | Bacteria | *Bacteroidota* | *Bacteroidia* | *Bacteroidales* | *Muribaculaceae* | *Muribaculaceae* |
| 4.060591511 | Bacteria | *Bacteroidota* | *Bacteroidia* | *Bacteroidales* | *Muribaculaceae* | *Muribaculaceae* |

**Supplementary Table 6: Taxonomic Markers Identified by LEfSe for Plant and Animal Diets**

| Feature | enrich _group | ef_lda | p-value | LDA_Adjusted |
| --- | --- | --- | --- | --- |
| *Muribaculaceae* | Plant | 5.233024635 | 5.40E-06 | 5.23302463 |
| *Allobaculum* | Plant | 4.64467002 | 1.88E-09 | 4.64467002 |
| *Bacteroides* | Plant | 4.565693781 | 7.05E-08 | 4.56569378 |
| *Escherichia-Shigella* | Plant | 4.379068769 | 7.89E-06 | 4.37906877 |
| *Parasutterella* | Plant | 4.22998311 | 4.69E-13 | 4.22998311 |
| *Tannerellaceae* | Plant | 4.188768412 | 1.77E-10 | 4.18876841 |
| *Lactobacillus* | Plant | 4.108421192 | 1.84E-05 | 4.10842119 |
| *Alloprevotella* | Plant | 4.032406516 | 3.72E-05 | 4.03240652 |
| *Muribaculum* | Plant | 3.875563718 | 1.80E-07 | 3.87556372 |
| *Atopobiaceae_g__* | Plant | 3.667431414 | 2.03E-06 | 3.66743141 |
| *Ileibacterium* | Plant | 3.66050386 | 2.48E-05 | 3.66050386 |
| *Parabacteroides* | Plant | 3.658075102 | 4.33E-08 | 3.6580751 |
| *Tannerellaceae_g__* | Plant | 3.637155508 | 3.96E-09 | 3.63715551 |
| *Mycoplasma* | Plant | 3.512558452 | 1.24E-13 | 3.51255845 |
| *Lachnospiraceae_UCG-001* | Plant | 3.486691951 | 1.06E-12 | 3.48669195 |
| *Helicobacter* | Plant | 3.430592519 | 1.74E-08 | 3.43059252 |
| *Erysipelatoclostridium* | Plant | 3.42541663 | 0.001001699 | 3.42541663 |
| *Gemella* | Plant | 3.404947204 | 1.49E-12 | 3.4049472 |
| *Clostridia_vadinBB60_group* | Plant | 3.349829721 | 0.000537424 | 3.34982972 |
| *Cetobacterium* | Plant | 3.345789548 | 5.29E-24 | 3.34578955 |
| *UCG-005* | Plant | 3.322054651 | 6.64E-13 | 3.32205465 |
| *Peptostreptococcaceae_g__* | Plant | 3.281207507 | 1.98E-06 | 3.28120751 |
| *Odoribacter* | Plant | 3.261282123 | 0.000978494 | 3.26128212 |
| *Corynebacteriaceae_g__* | Plant | 3.259661965 | 5.62E-07 | 3.25966196 |
| *Erysipelotrichaceae_g__* | Plant | 3.151128193 | 8.39E-07 | 3.15112819 |
| *A2* | Plant | 3.147567692 | 2.25E-05 | 3.14756769 |
| *Chloroplast* | Plant | 3.074413287 | 5.77E-20 | 3.07441329 |
| *Neisseriaceae_g__* | Plant | 3.064649391 | 1.38E-12 | 3.06464939 |
| *Butyricicoccus* | Plant | 3.054359327 | 8.76E-14 | 3.05435933 |
| *Prevotellaceae_UCG-001* | Plant | 3.047847732 | 5.63E-16 | 3.04784773 |
| *UCG-010* | Plant | 3.030970104 | 5.87E-06 | 3.0309701 |
| *Phascolarctobacterium* | Plant | 3.020812696 | 1.46E-17 | 3.0208127 |
| *Pseudomonas* | Plant | 3.00703619 | 1.25E-16 | 3.00703619 |
| *Rikenellaceae_g__* | Animal | 3.292743812 | 2.42E-07 | -3.2927438 |
| *Atopostipes* | Animal | 3.400705568 | 0.001739201 | -3.4007056 |
| *Intestinimonas* | Animal | 3.641121127 | 0.001865806 | -3.6411211 |
| *HT002* | Animal | 3.700185594 | 0.000483815 | -3.7001856 |
| *Clostridium_sensu_stricto_1* | Animal | 3.916651671 | 4.14E-06 | -3.9166517 |
| *Colidextribacter* | Animal | 4.038651745 | 0.007491941 | -4.0386517 |
| *Corynebacterium* | Animal | 4.436842966 | 0.008381161 | -4.436843 |
