## Supplementary material for "Dietary Protein Source Shapes Gut Microbial Structure and Predicted Function: A Meta-Analysis with Machine Learning": Meta_analysis Supplementary Figures

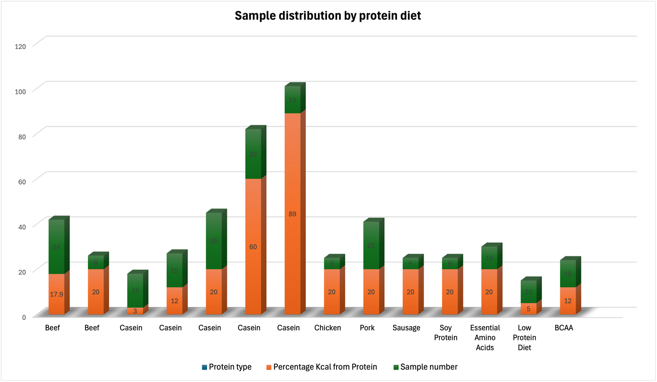

**1A**

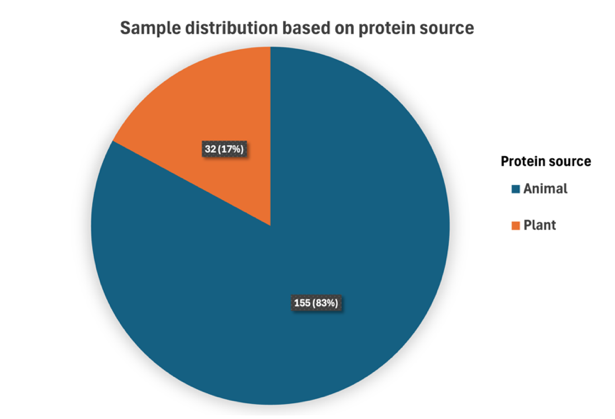

**1B**

**Supplementary Figure 1. Distribution of samples across included protein diets.**

**(A)** Sample distribution by specific protein types, showing protein type, percentage kcal from protein, and total sample number for each diet.

**(B)** Sample distribution based on protein source (animal vs. plant), illustrating the proportion of samples derived from animal-protein diets (83%) and plant-protein diets (17%).

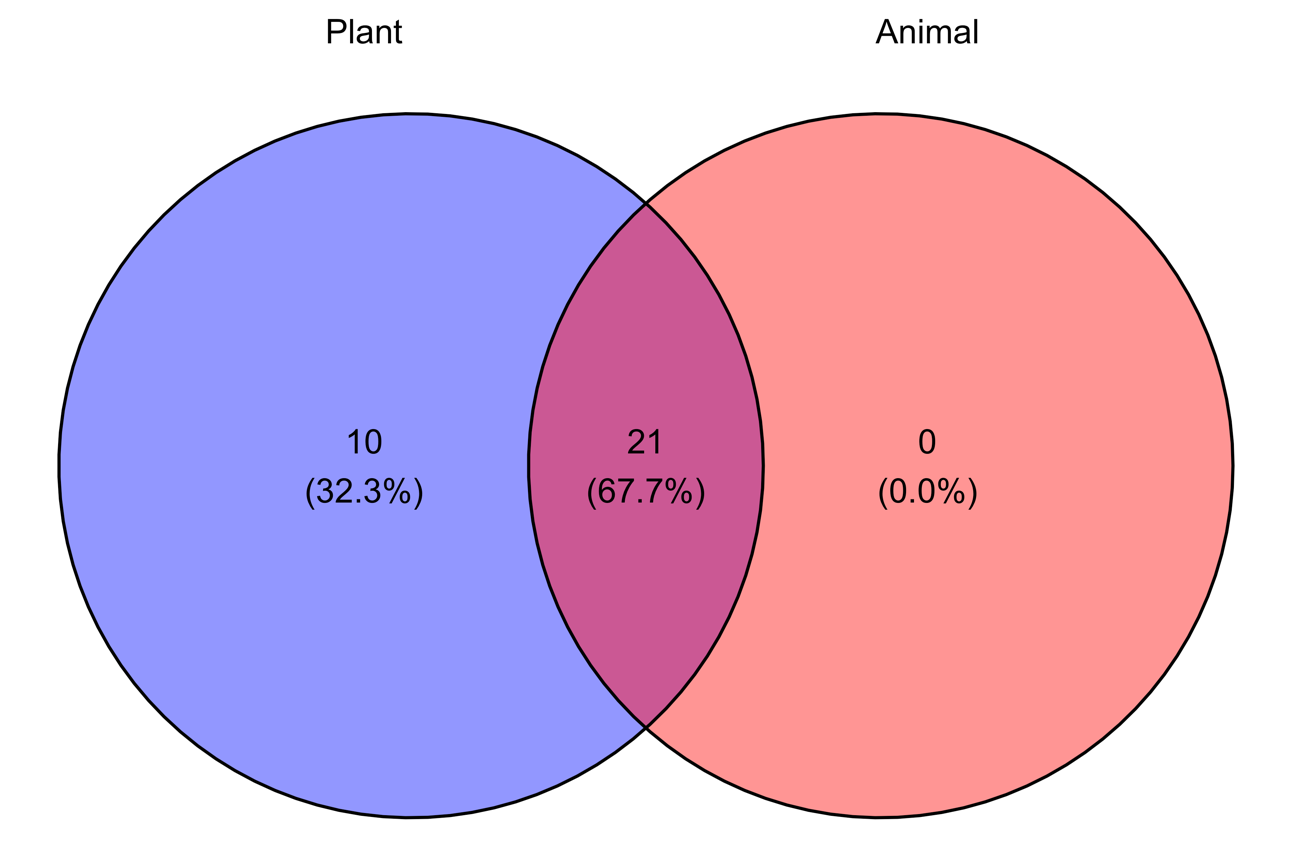

**2A**

**2B**

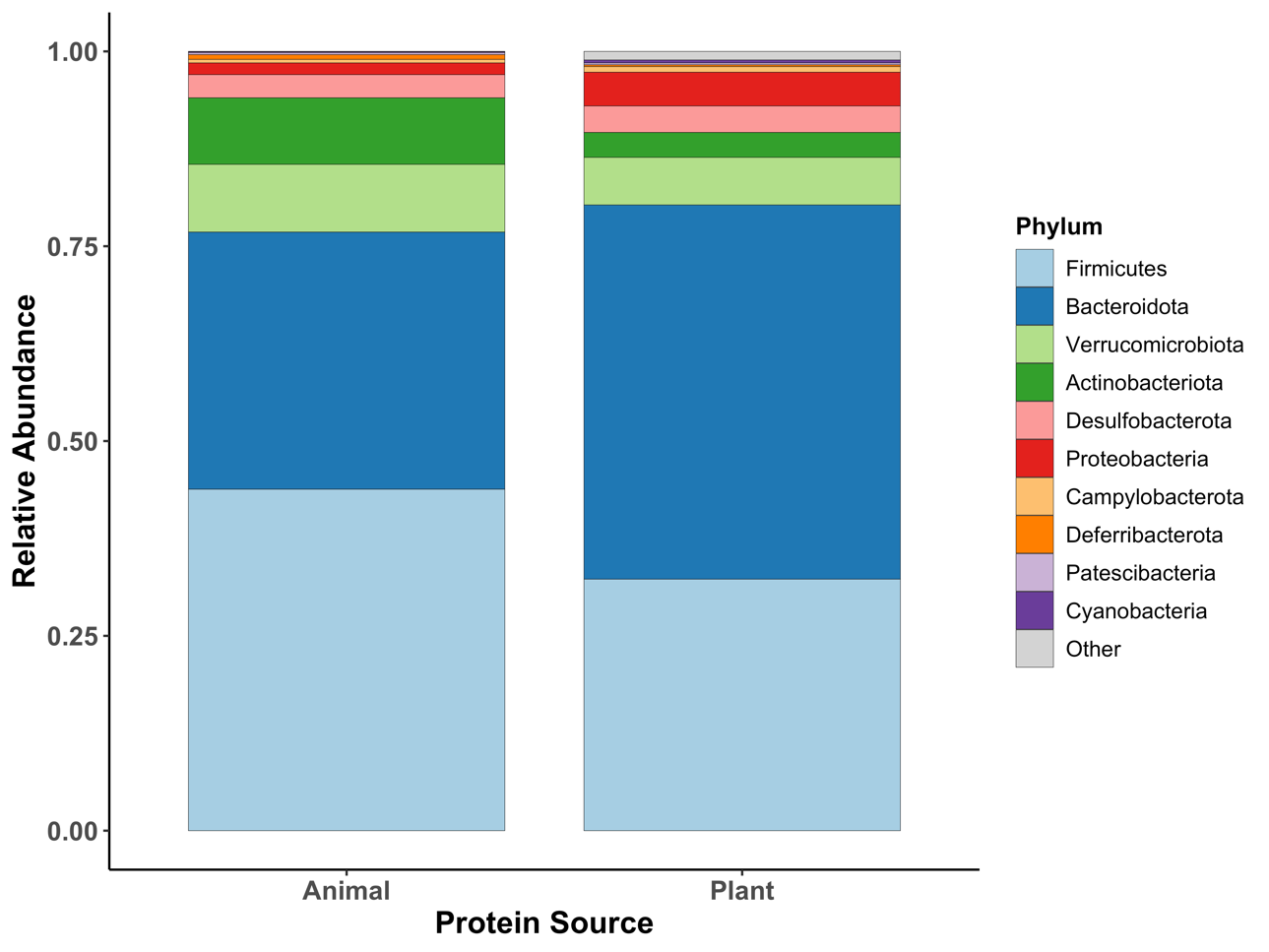

**Supplementary Figure 2. Core Microbiota Composition and Unique Microbial Taxa Across Protein Sources at the Phylum Level**

**(A)** Venn diagram showing the number and percentage of shared and unique microbial taxa between plant- and animal-protein diets at phylum level.

**(B)** Stacked bar plot depicting the relative abundance of dominant phyla across samples grouped by dietary protein source (animal vs. plant).

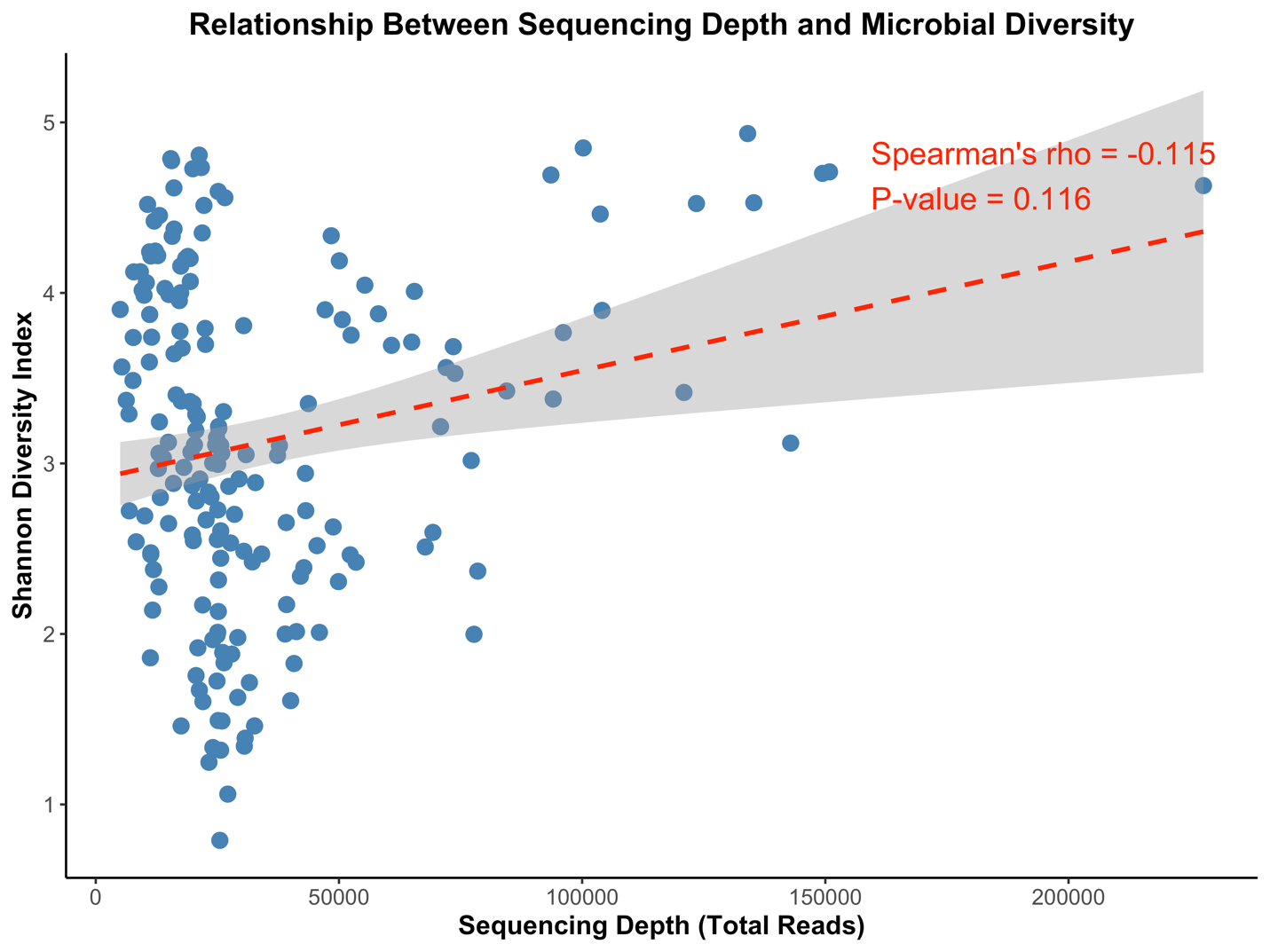

**3A**

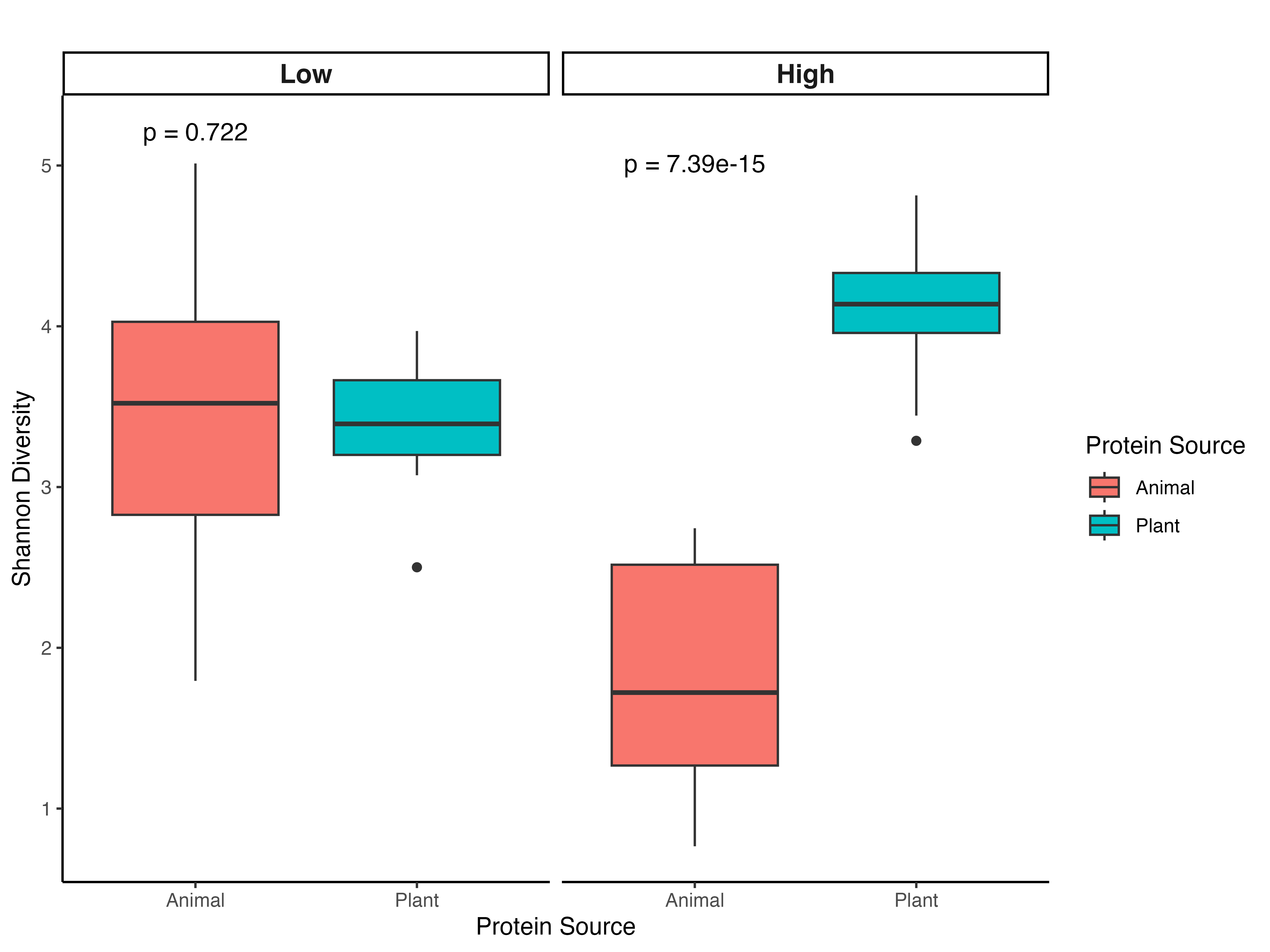

**3B**

**Supplementary Figure 3. Influence of Protein Concentration and Sequencing Depth on Alpha Diversity**

**A.** Relationship between sequencing depth and microbial alpha diversity (Shannon index). Each point represents an individual sample across all studies. Spearman correlation showed no significant association between sequencing depth and Shannon diversity (ρ = −0.115, p = 0.116), indicating that observed diversity differences were not driven by sequencing depth.

**B.** Interaction effect of protein source and concentration on gut microbial diversity (Shannon index). Boxplots show Shannon diversity across low and high protein concentrations for animal- and plant-based diets. Under high protein conditions, plant-based diets significantly increased microbial diversity compared to animal-based diets (p = 7.39 × 10⁻¹⁵). No significant difference was observed at low protein concentrations (p = 0.722).

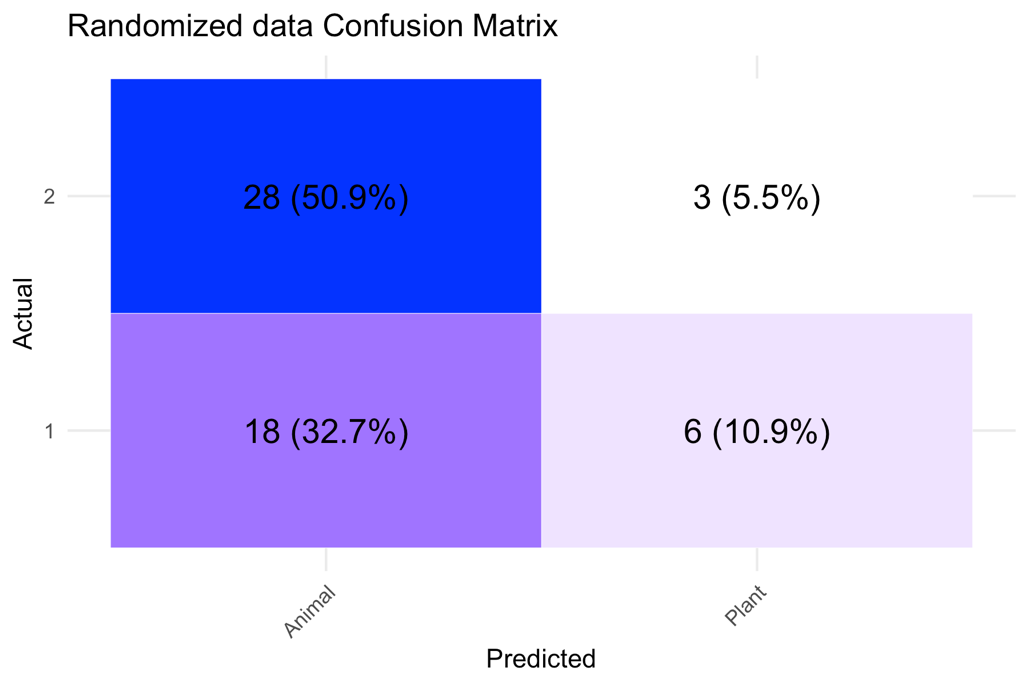

**4A**

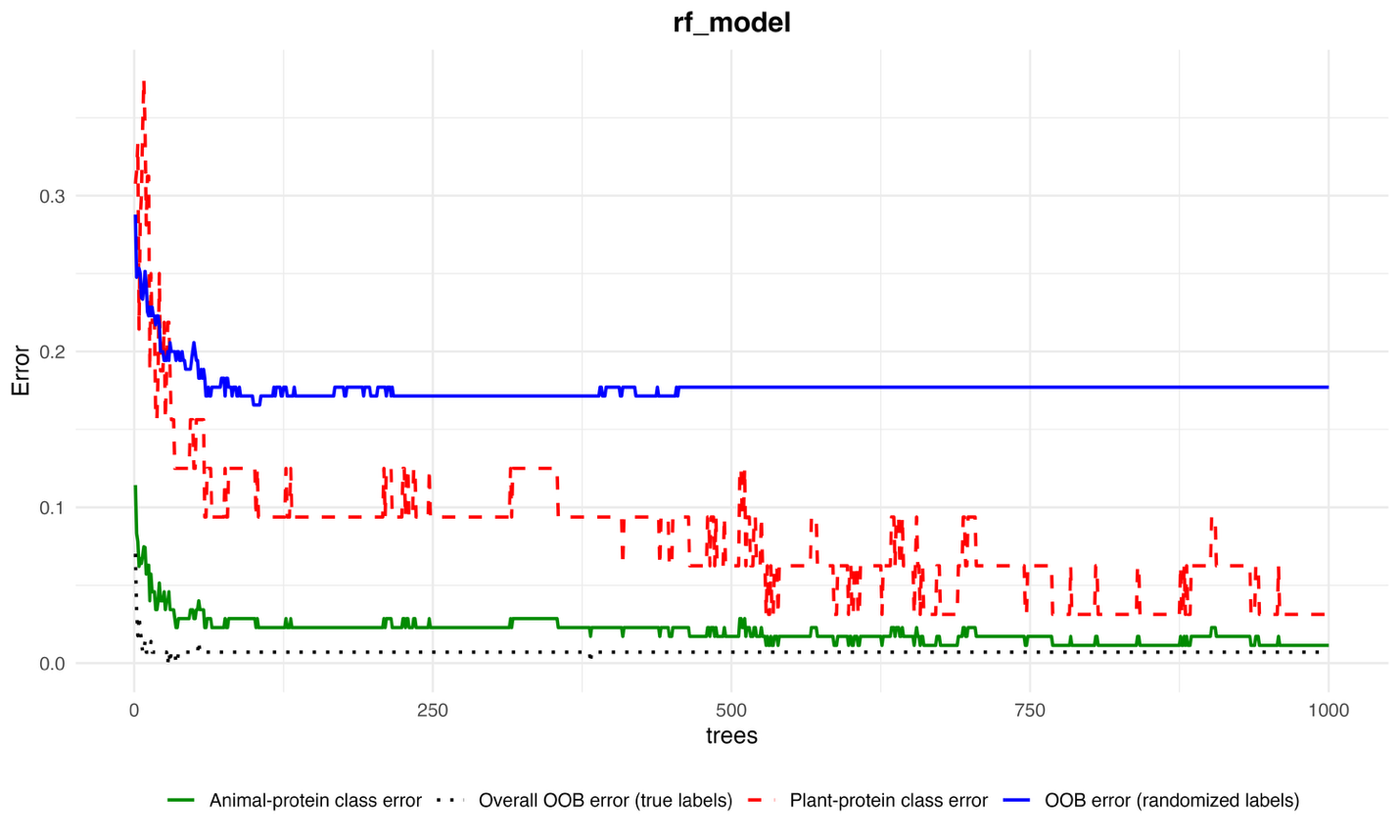

**4B**

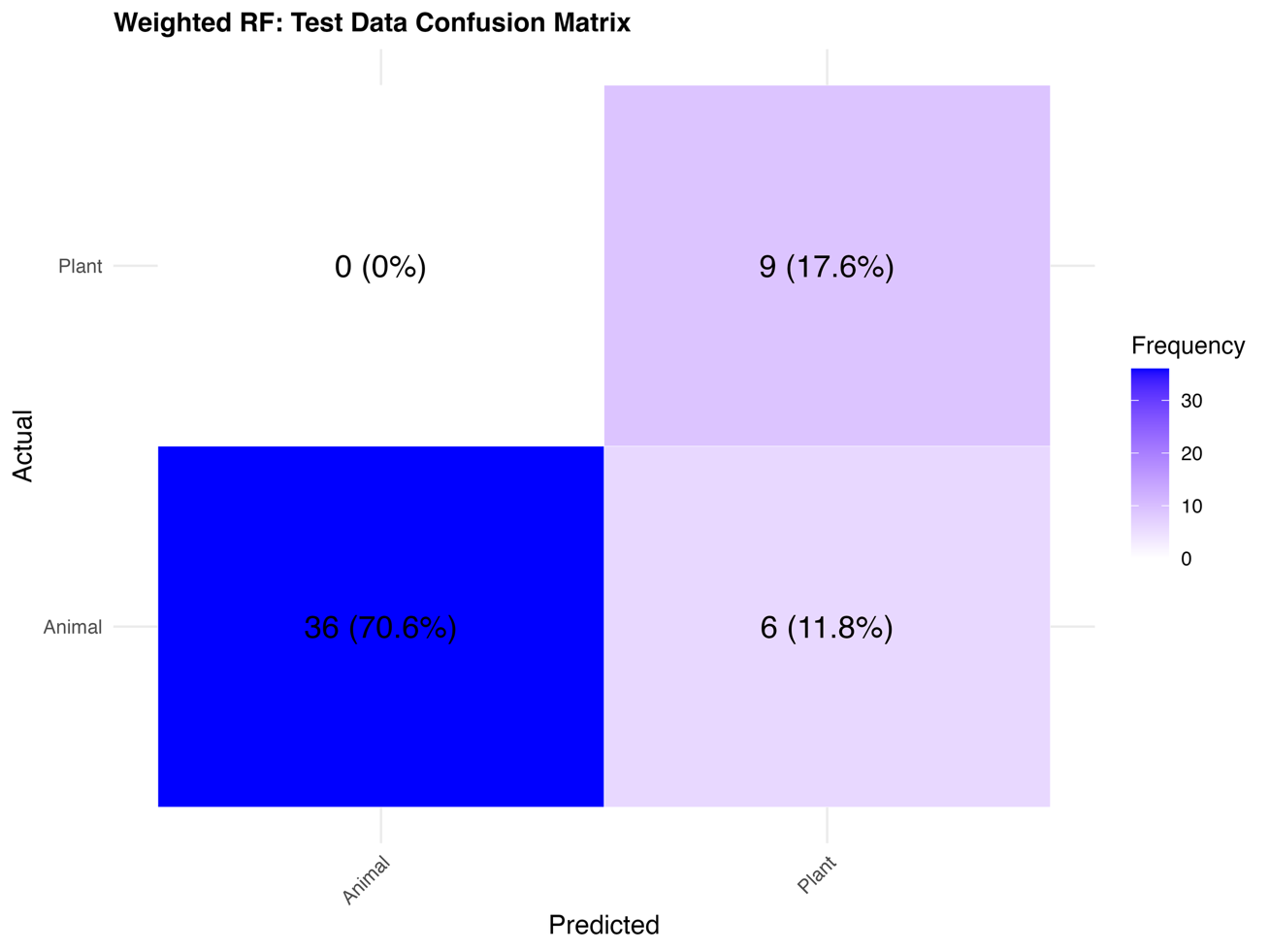

**4C**

**Supplementary Figure 4. Machine Learning Model Randomization and Error Diagnostics**

**(A)** Confusion matrix illustrating the classification performance of the randomized Random Forest model trained on shuffled protein-source labels. The poor accuracy demonstrates that the original model’s performance was not driven by random structure in the data.

**(B)** Error‐rate plot for the Random Forest model showing classification performance across 1,000 trees. The black dotted line represents the overall out-of-bag (OOB) error using the true class labels, the red dashed line shows the plant-protein class error, and the green solid line represents the animal-protein class error. The blue dot-dash line depicts the OOB error obtained after randomizing the class labels, serving as a baseline for model performance under noise. The rapid decline and subsequent stabilization of the true-label OOB and class-specific errors indicate model convergence and robust discriminatory power relative to the randomized baseline.

**(C)** Confusion matrix of the class-weighted Random Forest model showing predictions for the test dataset. The model achieves high discrimination between plant- and animal-based diets compared with randomized data, confirming true signal.

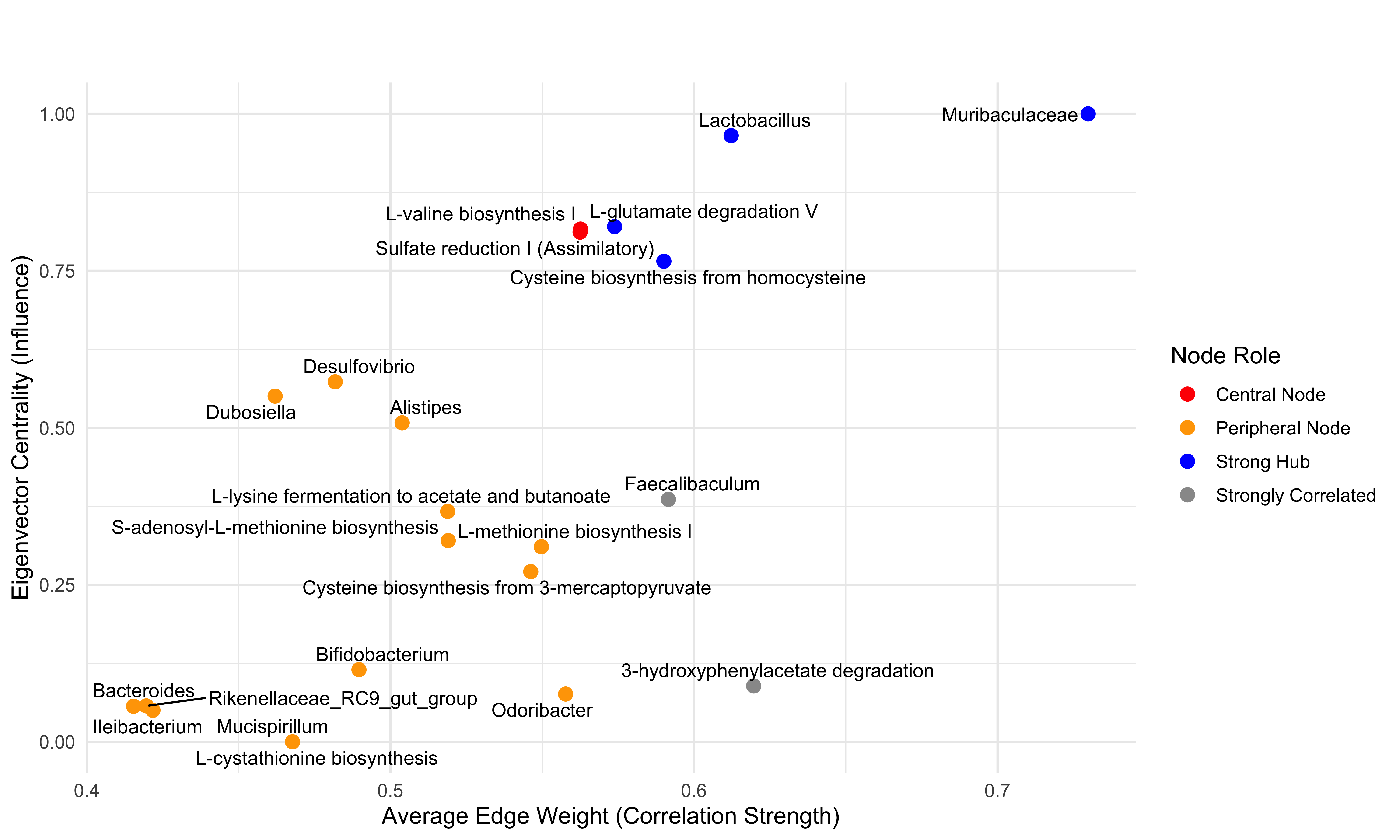

**5**

**Supplementary Figure 5. Network centrality plot of microbial taxa and predicted functional pathways.**

This scatterplot displays node-level topological properties within the taxon–function correlation network. The x-axis represents average edge weight (correlation strength), and the y-axis indicates eigenvector centrality (node influence). Node color reflects functional role: blue nodes are strong hubs with high centrality and correlation; red nodes are central connectors; orange nodes are peripheral; and grey nodes are strongly correlated but less influential
